# RNAi-based screen for pigmentation in *Drosophila melanogaster* reveals regulators of brain dopamine and sleep

**DOI:** 10.1101/2023.07.20.549932

**Authors:** Samantha L Deal, Danqing Bei, Shelley B Gibson, Harim Delgado-Seo, Yoko Fujita, Kyla Wilwayco, Elaine S Seto, Amita Sehgal, Shinya Yamamoto

## Abstract

The dopaminergic system has been extensively studied for its role in behavior and neurological diseases. Despite this, we still know little about how dopamine levels are regulated *in vivo*. To identify regulators of dopamine, we utilized *Drosophila melanogaster* cuticle pigmentation as a readout, where dopamine is used as a precursor to melanin. We started by measuring dopamine from known pigmentation mutants (e.g. *tan*, *ebony*, *black*) and then performed an RNAi-based screen to identify new regulators. We found 153 hits, which were enriched for developmental signaling pathways and mitochondria-associated proteins. From 35 prioritized candidates, 11 had an effect on head dopamine levels. Effects on brain dopamine were mild even when the rate-limiting synthesis enzyme *Tyrosine hydroxylase (TH)* was knocked down, suggesting changes in dopamine levels are tightly regulated in the nervous system. We pursued two of our hits that reduced brain dopamine levels, *clueless* and *mask*. Further examination suggests that *mask* regulates transcription of *TH* and affects dopamine-dependent sleep patterns. In summary, by studying genes that affect cuticle pigmentation, we were able to identify genes that affect dopamine metabolism as well as a novel regulator of behavior.

## Introduction

Dopamine is a conserved neurotransmitter that functions in the nervous system to regulate a variety of behaviors, including learning & memory, locomotion, reward, and sleep. In humans, disruptions in dopamine have been associated with neurological and neuropsychiatric disorders, including addiction, depression, sleep disorders, and schizophrenia.^1–3^ In addition, one of the hallmarks of Parkinson’s disease is early degeneration of dopaminergic neurons, and Parkinson’s patients are often given a dopamine precursor, l-dihydroxyphenylalanine (L-DOPA), as a method for treating symptoms.^4,5^ Most genetic research on dopamine biology has focused on dopaminergic signaling and transport. However, changes in dopamine levels can have a great impact on behavioral outcomes.^6–8^ Thus, identifying genes that affect dopamine levels is critical to understanding and potentially treating dopamine-associated disorders.

Dopamine synthesis is a conserved process, whereby the amino acid tyrosine is converted into L-DOPA via the rate-limiting enzyme, Tyrosine hydroxylase (TH).^9^ Then, L-DOPA is converted into dopamine via the enzyme Dopa decarboxylase (Ddc).^10^ In mammals, dopamine is degraded via oxidation or methylation using the enzymes Monoamine oxidase (MAO) and Catechol-O-methyltransferase (COMT).^11^ In invertebrates, dopamine degradation goes through β-alanylation via the β-alanyl amine synthase (encoded by the *ebony* gene in *Drosophila melanogaster*) and acetylation via the acetyltransferase (encoded by the *speck* gene in *Drosophila*).^12^ In addition, insects also convert dopamine into melanin to form proper cuticle pigment and structure. ^13–15^For example, knockdown of *TH* (also known as *pale (ple)* in *Drosophila*) or *Ddc* leads to a paler cuticle, and reduction of *ebony* or *speck* leads to a darker cuticle.^13–15^

Historically, *Drosophila* has been used to study dopamine biology in multiple contexts. In particular, the fly has been instrumental in dissecting the role of dopamine in learning and memory.^16,17^ Moreover, an extensive genetic toolkit has made it a great model system for dissecting dopaminergic neural circuitry and signaling.^18–20^ Lastly, *Drosophila* has been used to study dopamine-related neurological diseases. In particular, the fly was instrumental in identifying and investigating the cellular functions of Parkinson disease’s genetic risk factors, *PINK1* and *PRKN* (i.e. *Pink1* and *parkin* in ^21^*Drosophila*).^21^

As well as being a tool for dissecting neural circuitry, the fast generation time and low maintenance cost makes *Drosophila* a great system for large-scale screening. Previously, a RNAi screen (11,619 genes, ∼89% of genome) was performed to identify genes that affect mechanosensory organs and cuticle formation in the *Drosophila* thorax.^22^ In this screen, the researchers found 458 genes that affected cuticle color upon knockdown, but no further validation was performed.

Here, we utilize the fly cuticle to identify novel regulators of dopamine. We start by systematically examining classical pigmentation genes for their effect on dopamine in the head and brain. These experiments revealed that genes involved in dopamine synthesis cause the expected reduction in dopamine. On the other hand, genes involved in dopamine degradation either have no effect or unexpectedly show reduced dopamine. We go on to characterize 458 genes identified from the RNAi screen by Mummery-Widmer et al. (2009). We tested 330 of them with independent RNAi lines and validated 153 genes (∼46%) with consistent cuticle pigmentation defects. We go on to examine 35 prioritized gene hits for their effect on dopamine levels. From this analysis, we found two genes, *mask* and *clu*, that reduced brain dopamine levels upon knockdown in dopaminergic neurons. Lastly, we used molecular biology and sleep behavioral studies to show that *mask* appears to alter brain dopamine through regulation of *TH* transcription.

## Results

### *Drosophila* pigmentation genes affect dopamine in unexpected ways

To systematically test if changes in pigmentation genes affect dopamine as theorized in the field, we characterized loss-of-function alleles of genes involved in dopamine metabolism (Figure 1A) for pigmentation phenotypes and dopamine levels. Since the enzymes required for dopamine synthesis (TH and Ddc) are essential for survival,^23,24^ we took advantage of established RNAi lines and UAS/GAL4 system.^25–28^ It is known that reduction of *TH* or *Ddc* causes pale pigmentation and reduced dopamine.^9,10,14^ We validated our method by knocking down *TH* in the middle thorax using *pnr-GAL4* along with a verified RNAi,^29^ which causes a strong pale cuticle phenotype (Figures 1B and 1C). We measured dopamine using HPLC, and in our hands, this RNAi reduces head dopamine levels by ∼60% (Figure 1E) and brain dopamine levels by ∼32% (Figure 1E’). *Ddc* knockdown causes a weaker effect than *TH* RNAi in the cuticle (Figures 1B and 1D) and the head (∼35% dopamine reduction, Figure 1E) and has no significant effect on dopamine in the brain (Figure 1E’). This may be due to the strength of the RNAi and/or enzyme level requirements (since TH is the rate-limiting enzyme).

**Figure 1.**
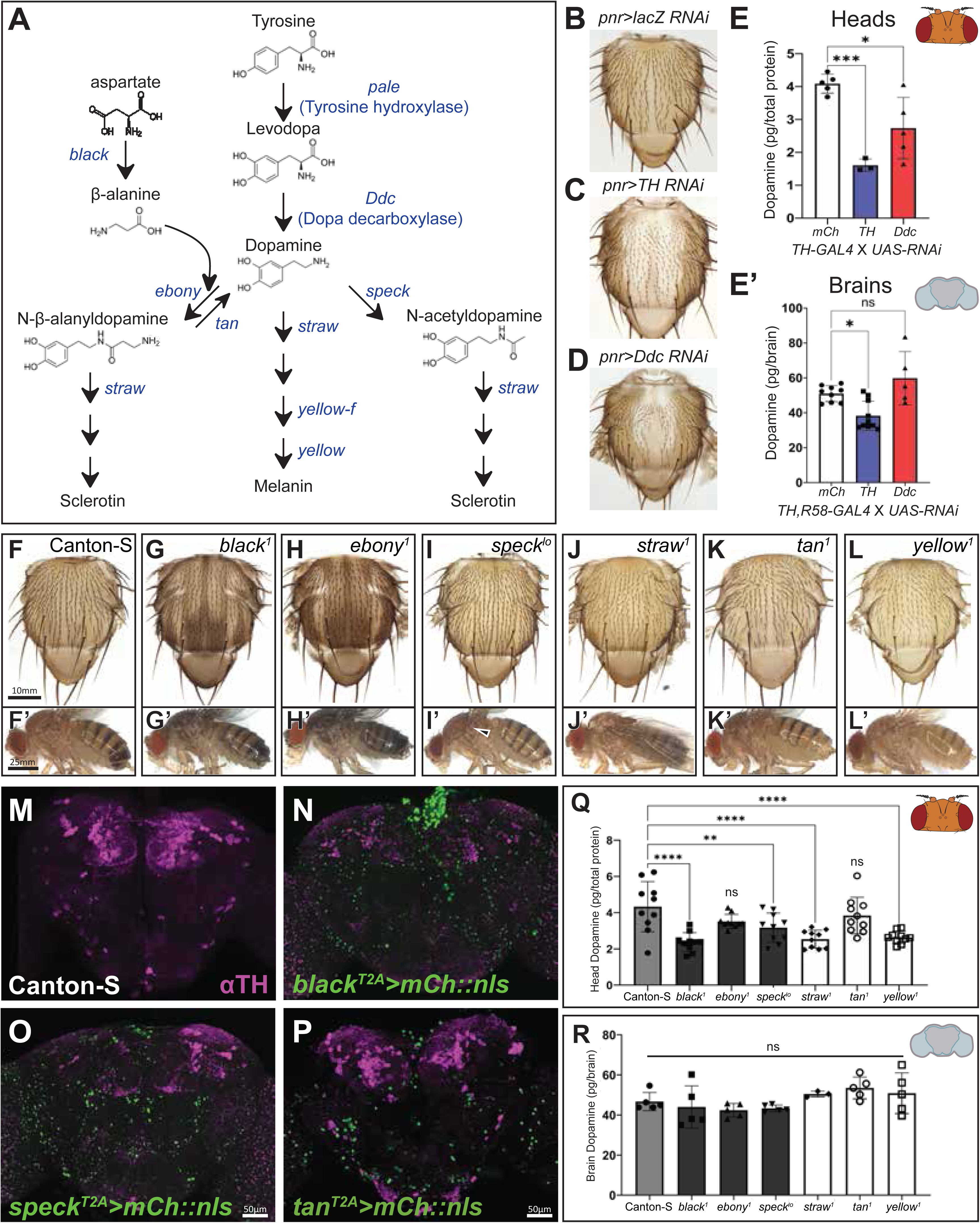
Manipulation of pigmentation enzymes have unexpected effects on dopamine. (A) *Drosophila* dopamine metabolism pathway. Gene names encoding pigmentation enzymes are in blue and metabolites are in black. (B-D) Thorax phenotypes observed upon knockdown of Tyrosine hydroxylase (TH, also known as Pale) (C) and Dopa decarboxylase (Ddc) (D) compared to control (B). (E-E’) Dopamine levels, measured using HPLC, from the heads (E) and brains (E’) upon knockdown of *TH* and *Ddc*. Pigmentation phenotypes observed mutant alleles in the thorax (F-L) and the body (F’-L’) (Arrow in I’ indicates the darker region characteristic of *speck* mutants). *T2A-GAL4* alleles for *black*, *speck*, and *tan* adult brain expression with dopaminergic neurons (DAN) staining in magenta (DANs) (M-P). (Q) Dopamine, measured using HPLC, in the heads of classical pigmentation mutants, *black^1^*, *ebony^1^*, *straw^1^*, and *yellow^1^*. (R) Dopamine levels in dissected brains of classical pigmentation mutants. Ordinary one-way ANOVA test with Dunlett multiple comparisons test. *=p<0.05, **=p<0.01, ***=p<0.001, ****=p<0.0001. Error bars show standard deviation (SD).

To test genes implicated in dopamine degradation and cuticle development (i.e. pigmentation and structural rigidity), we examined known mutant alleles. These alleles cause darker or paler cuticles (Figures 1F-1L and 1F’-1L’) depending on their role in melanin metabolism. Unexpectedly, *black*, *speck*, *straw* and *yellow* mutants reduce head dopamine levels (measured using HPLC, Figure 1Q), while *ebony* and *tan* mutants had no significant effect (Figure 1Q). Next, we checked the brain expression patterns of pigmentation enzymes. Based on our own expression analysis (Figures 1M-1P) and previously reported expression data,^30,31^ most are expressed in the fly brain, but their expression does not appear to overlap with DANs. None of these pigmentation gene mutations had a significant effect on brain dopamine (Figure 1R).

Together this data shows that changes in dopamine synthesis cause a paler cuticle, which reflects a reduction in dopamine, but changes in dopamine metabolism enzymes downstream of dopamine synthesis have pigmentation phenotypes that do not correlate with brain dopamine level differences.

### RNAi-based cuticle pigmentation screen

Mining a previously published genome-wide scale RNAi screen,^22^ we identified 458 genes with pigmentation defects. To check if these phenotypes can be reproduced, we located 718 additional UAS-RNAi lines corresponding to 330 genes. 426 lines came from the National Institute of Genetics in Mishima, Japan,^32^ and 292 came from the Transgenic RNAi Project (TRiP) at Harvard Medical School.^25^ 153 genes caused pigmentation defects in at least one additional RNAi line, which validates the results for 46.4% of the previous screen’s RNAi lines (VDRC collection, Figure 2A).

**Figure 2.**
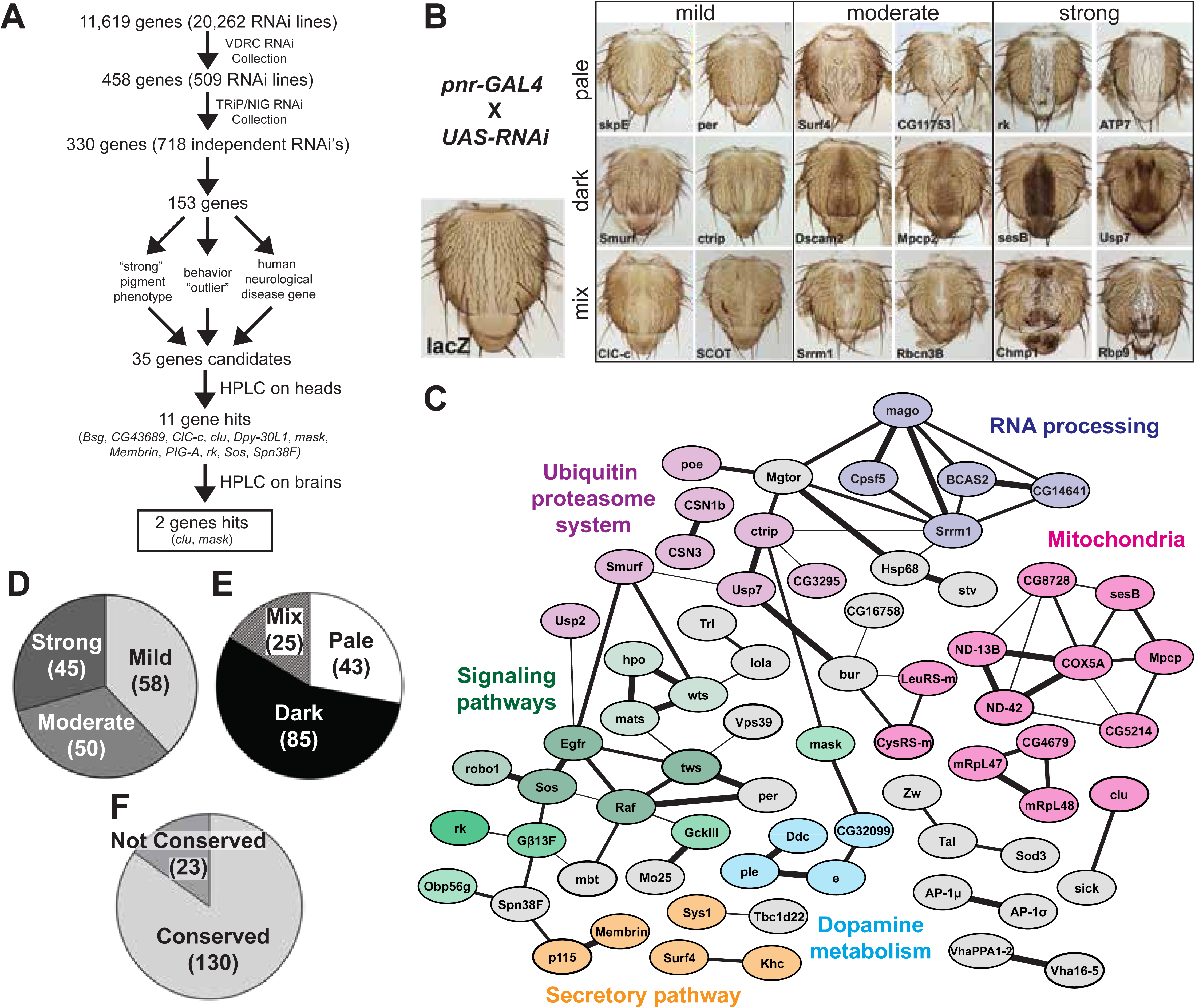
RNAi pigmentation screen reveals a broad spectrum of phenotypes and classes of genes. (A) Flow chart depicting screening process. (B) Examples of the phenotypes observed from the RNAi-based pigmentation screen. (C) Protein-protein interaction network for 153 high confidence pigmentation gene hits from the RNAi-screen using STRING network analysis. Strength of line indicates confidence or strength of data support. Genes with no known interactions are not displayed. (D-F) Pie charts for the 153 pigmentation gene hits, including strength of phenotype (D), phenotype color (E), and conservation (F).

Our RNAi-based secondary screen produced a spectrum of phenotypes (Figure 2B), but the phenotypes strengths (mild, moderate, or strong) were equally represented (mild = 38%, moderate = 33%, strong = 29%, Figure 2D). Pigmentation color changes were classified in three primary categories, pale, dark, or mix (a combination of pale and dark in a single thorax), which were not equally represented (Figure 2E). Dark cuticle phenotypes were the most common (56%), with pale and mixed cuticle phenotypes seen in 28% and 16% of genes, respectively.

We performed a protein-network analysis (Figure 2C) and found a group of proteins previously linked to cuticle pigmentation as well as clusters related to RNA processing, ubiquitin proteasome system, mitochondria-associated proteins, and signaling pathway genes, Hippo and EGF (epidermal growth factor) signaling in particular. GO enrichment analysis highlighted terms associated with cuticle pigmentation, protein stability, and several signaling pathways (Figure S1). We also observed that 85% of our genes were conserved between *Drosophila* and human, an enrichment from the 68% of conserved genes in the original screen gene list (11, 619, Figures 2F and S2).

In summary, the cuticle pigmentation screen identified a spectrum of pale, dark and mixed phenotypes in 153 genes, showing a validation of ∼50% from the original screen. Close examination of these hits highlights conserved molecular processes.

### Many pigmentation genes isolated have a neurological consequence

To investigate if our gene candidates have a neurological consequence, we first checked if their homologs are associated with a neurological disease. Importantly, we defined homologs or conserved genes as genes where an ortholog prediction algorithm (DRSC Integrative Ortholog Prediction Tool) had a human gene with a score greater than or equal to three. We found that 51.2% of the fly gene homologs are associated with a disease, 78% of those have at least one disease with neurological phenotypes (Table 1 and Figure S2), such as intellectual disability, autism spectrum disorder, ataxia, epilepsy, and cerebral atrophy.

**Table 1.**
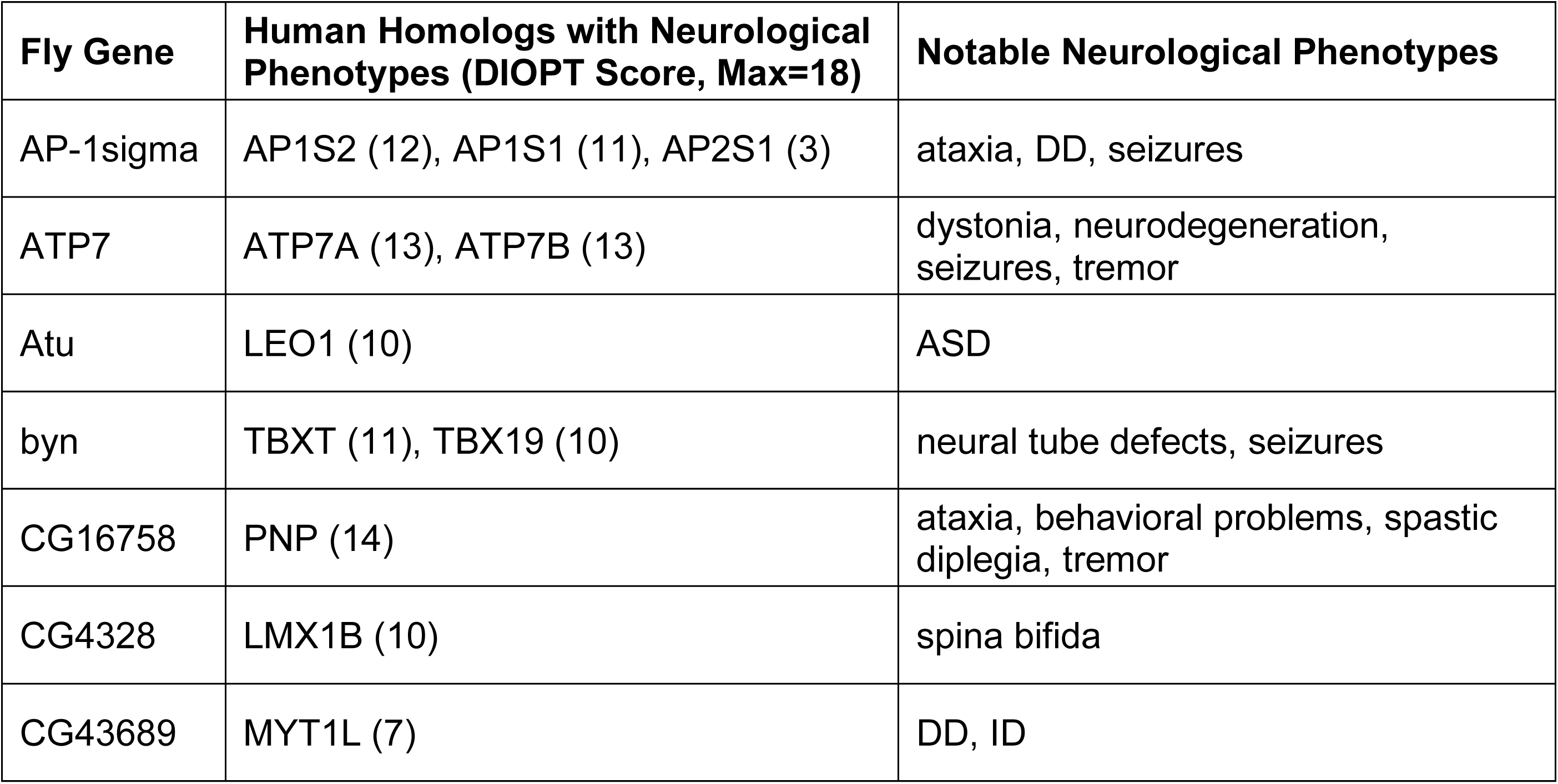
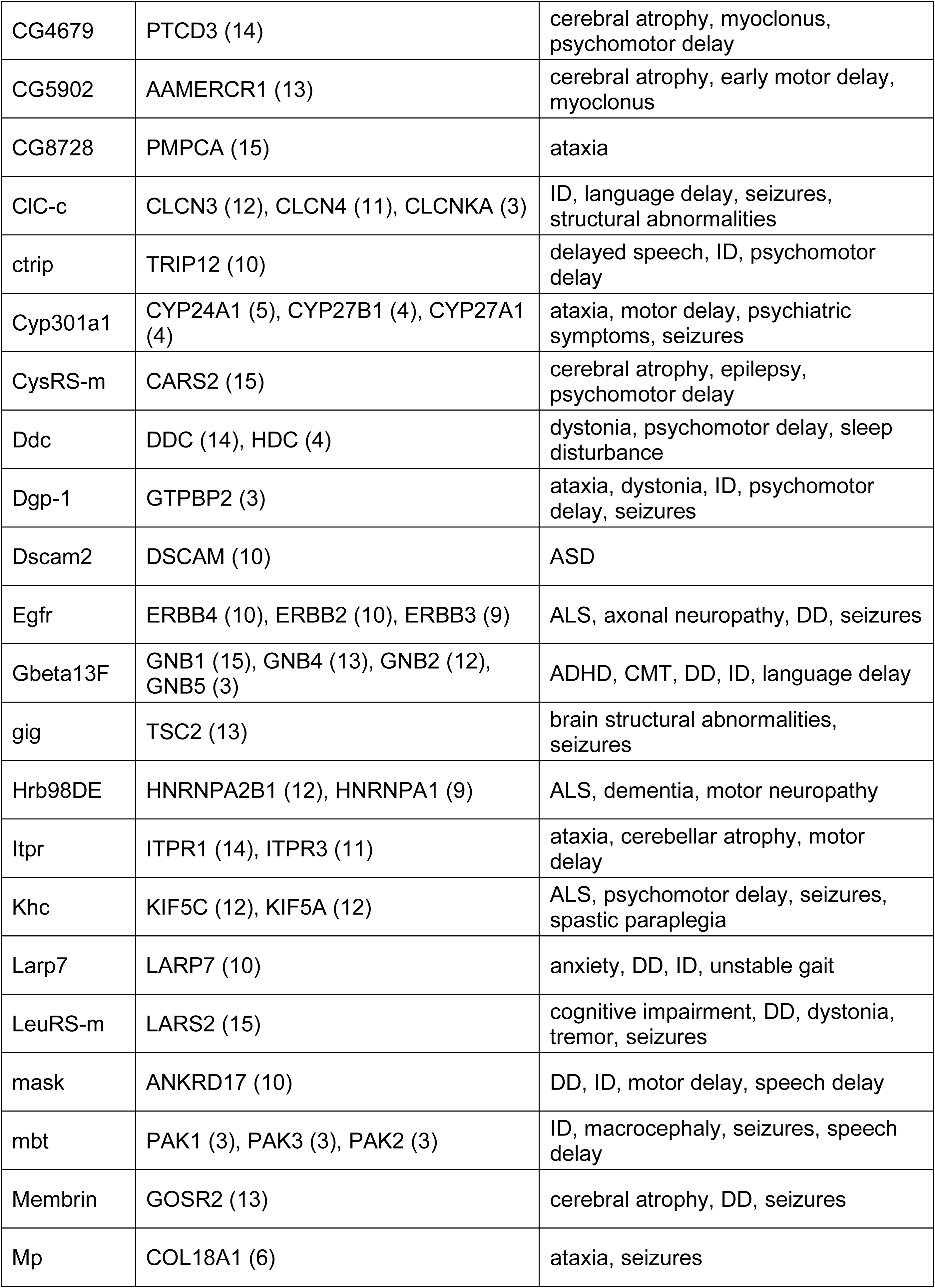
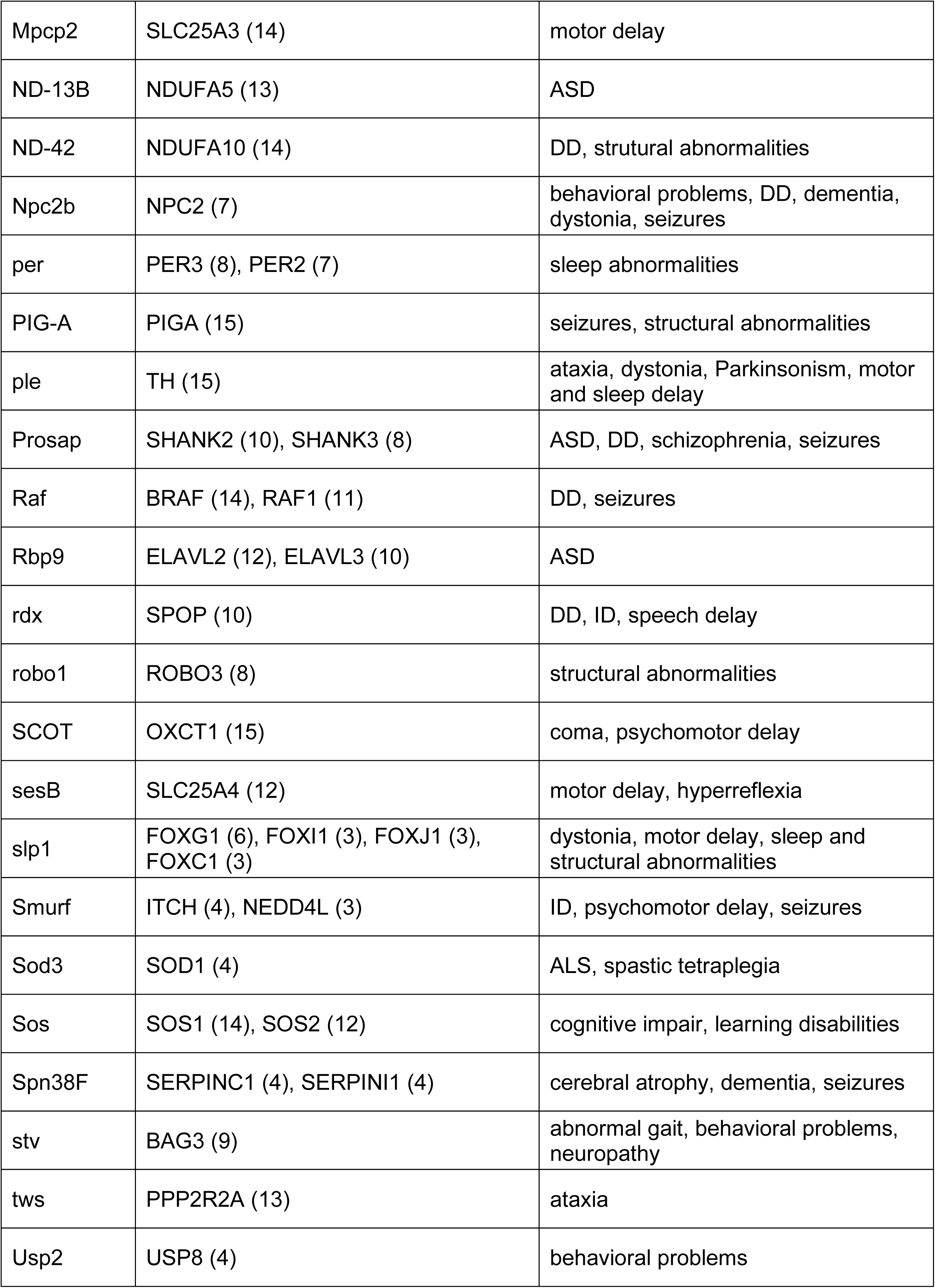
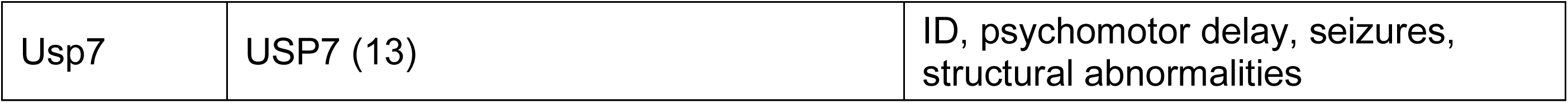
Fly pigmentation genes identified from this screen with neurological disease associated homologs in human. Homologs were defined as a Drosophila Integrative Orthology Prediction Tool (DIOPT, version 8.0) score greater than or equal to three. Neurological disease associations were based on Online Mendelian Inheritance in Man (OMIM)_clincal disease phenotypes and high-confident ASD genes listed in Simons Foundation Autism Research Initiative (SFARI) gene database (class S, 1, 2 or 3). ASD=Autism Spectrum Disorder, DD=Developmental Delay, ID=Intellectual Disability, ALS=Amytrophic Lateral Sclerosis, CMT=Charcot-Marie-Tooth disease.

Since dopamine is known to regulate locomotion and sleep,^22,33^ we screened for locomotion and sleep defects using the Drosophila Activity Monitor (DAM).^34^ We tested 274 RNAi lines from the NIG, TRiP, and VDRC RNAi collections corresponding to the validated 153 gene hits.^35–37^ We scored the lines for total locomotion, total sleep, sleep bout length, and sleep latency during the night and day (Figure 3). There was a broad range in the RNAi lines for each of these phenotypes (Figure 3C). We classified RNAi lines as “outliers” if they were two standard deviations from the mean for a given phenotype (locomotion: 23 lines, total sleep: 20 lines, sleep bout length: 21 lines, sleep latency: 25 lines). While total sleep and locomotion strongly overlapped with each other, sleep latency and sleep bout length did not overlap with the other sleep phenotypes (Figure 3D). We assessed if certain gene categories (>10 genes) showed any distinct behavioral patterns (Figure 2C). There was no change in total locomotion or total sleep in the categories assessed (Figures 3F and 3G). However, each category showed a distinct pattern in sleep bout length and latency (Figures 3F and 3H). This could indicate a distinction in how different gene functions could affect sleep. Since genes that affect sleep can often show distinct phenotypes (e.g. total sleep, bout number, latency, etc.) at distinct times of day,^38^ any gene with one phenotype during the day or night was classified as an “outlier”. This produced 60 lines, corresponding to 50 genes.

**Figure 3.**
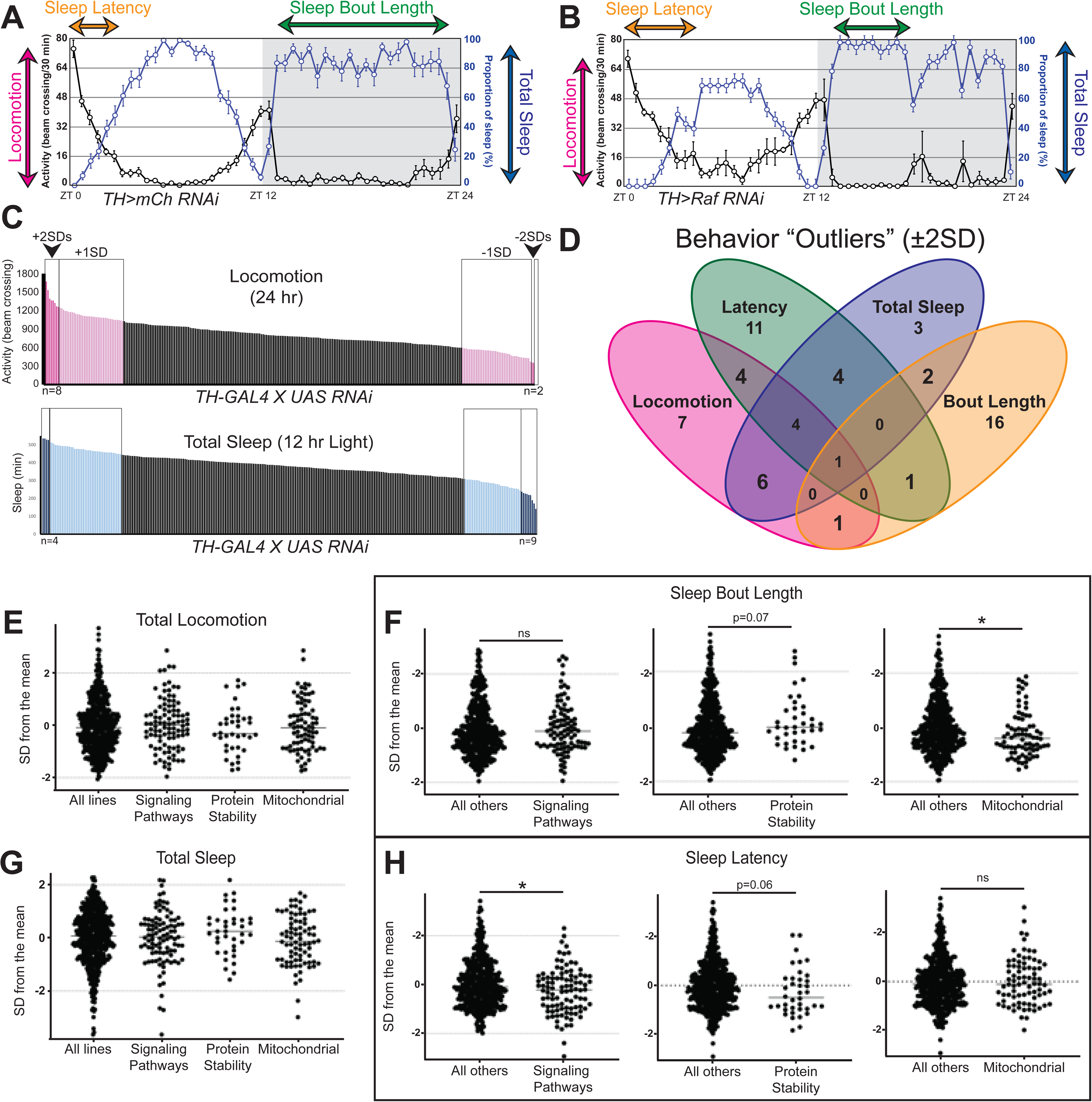
***Drosophila* Activity Monitor (DAM) screen identifies behavior “outliers” and highlights distinct phenotypes for gene categories.** (A-B) Representative activity and sleep data measured in a neutral RNAi line (*mCherry RNAi*, A) and a behavioral “outlier” (*Raf RNAi*, B) knocked down in DANs (*TH-GAL4*). (C) Bar graph showing all RNAi lines tested for 24hr locomotion and total sleep during the 12hr light period, where light pink/blue represents fly lines ±1SD away from the mean, and dark pink/blue are fly lines ±2SDs away from the mean (a.k.a. “outliers”). (D) Venn diagram of “outliers” for the four phenotypes examined (sleep latency, sleep bout length, locomotion, and total sleep). (E-H) Behavioral analysis of lines in three gene categories captured in the pigmentation screen, including locomotion (E), sleep latency (F), sleep bout duration (G), and total sleep (H). Individual t-tests were performed to assess if genes in a category were significantly different to all other genes tested, and only those with a p<0.10 are reported, with *=p<0.05.

Taken together, 77 of our 132 conserved genes are likely to have a neurological function, because they are implicated in human neurological disease (52 conserved genes) or were classified as a behavioral “outlier” (45 conserved genes).

### 11 genes alter dopamine levels in the fly head

To prioritize our gene list for dopamine measurement, we classified which of our genes were conserved in humans, as previously noted (132 conserved/153 pigmentation genes). Then, we prioritized these based on whether knockdown of these genes showed a strong pigmentation effect (38 genes), a behavioral effect in the DAM (45 genes), and/or a human neurological disease association (52 genes). 72% (95/132 conserved genes) were represented in one of these categories, and 29% (39/132) were represented in at least two (Figures 4A). We classified genes in two or more categories as priority hits and tested them for changes in dopamine using High Performance Liquid Chromatography (HPLC). We repeated previous results showing that knocking down *TH* or overexpressing a *TH* cDNA using *TH-GAL4* significantly reduces or increases head dopamine without effecting serotonin (Figures 4B and 4C).^29^

**Figure 4.**
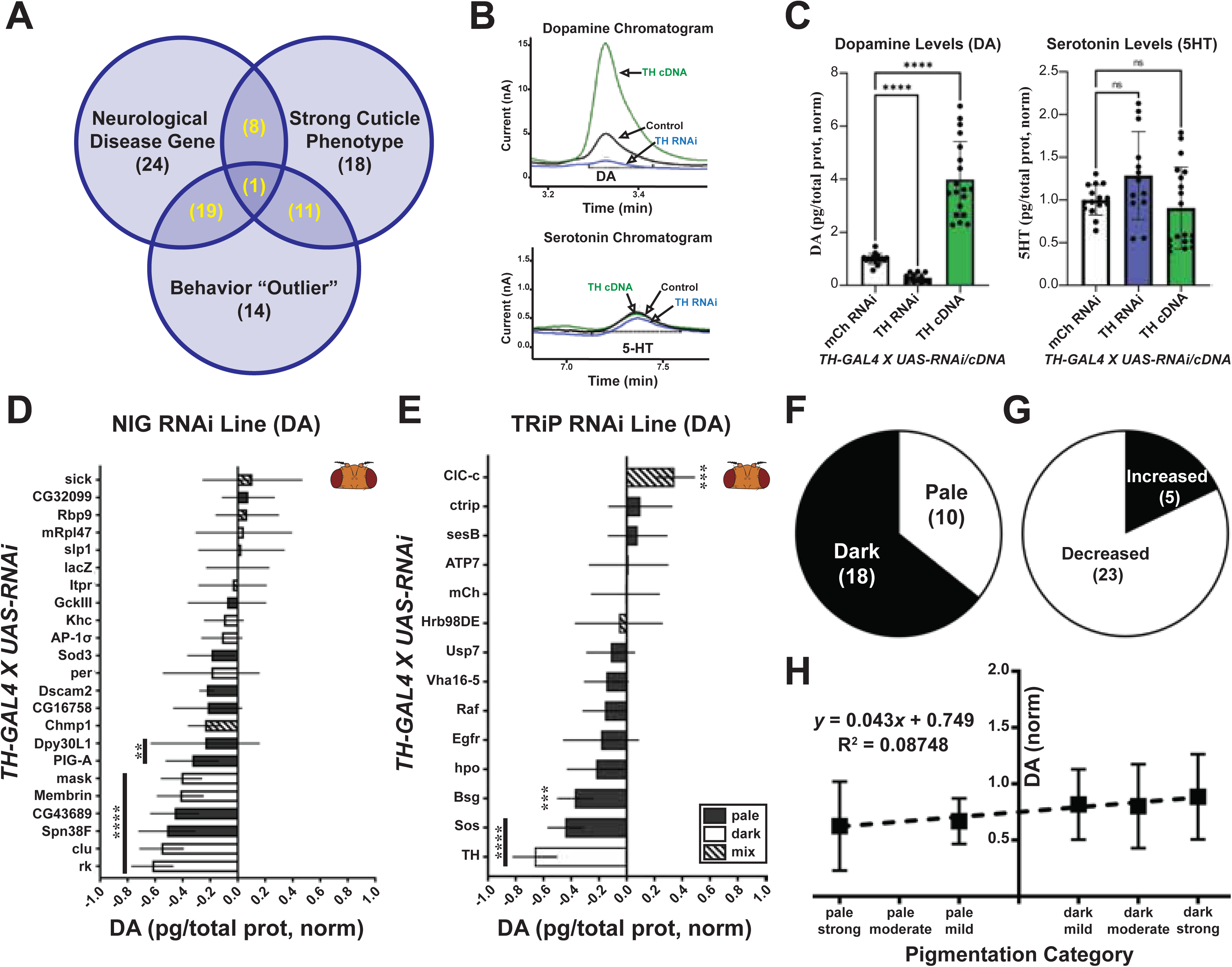
Eleven genes alter head dopamine upon knockdown. (A) Venn diagram of the categories used to prioritize 39 genes (in yellow). (B) Representative chromatograms for dopamine (DA) and serotonin (5-HT) upon knockdown or overexpression of TH in Dopaminergic cells (*TH-GAL4*). (C) Quantification of DA and 5-HT from chromatograms. (D-E) HPLC analysis on prioritized genes for UAS-RNAi lines from NIG (D) and TRiP (E). (F-G) Pie charts for all genes for cuticle color (F) and trend in dopamine (G). (H) XY plot comparing pigmentation phenotype to dopamine levels. For (C) Brown-Forsythe ANOVA test performed with Dunnett’s T3 multiple comparisons test. For (D) and (E) ordinary One-way ANOVA test with Dunnett’s multiple comparisons test. **=p<0.01, ***=p<0.001, ****=p<0.0001. Error bars represent SD.

We found that 11 of 35 prioritized genes (note: some not tested due to lethality or stock issues) significantly altered head dopamine levels (Figures 4D and 4E). Since control NIG lines showed about ∼80% dopamine levels compared to the TRiP collection (Figure S3), we compared the DA levels to control lines appropriate for each collection (e.g. UAS-lacZ RNAi). Unlike cuticle pigmentation, where 65% of genes caused a dark cuticle (excludes mixed cuticle phenotype), most genes trended toward a reduction in dopamine (71%, 25/35, Figures 4F and 4G). There was no correlation between the cuticle color and the dopamine level in the fly head (Figure 4H).

In summary, our HPLC analysis on fly heads revealed 11 genes that significantly affect dopamine levels. We saw a trend in reduction in dopamine across all lines tested, and we observed no correlation between cuticle color and dopamine level.

### Brain studies reveal *mask* and *clueless* as regulators of brain dopamine

To determine which of the 11 gene hits have consequences in the brain, we examined if their expression ^39–41^overlapped with DANs. Looking at single-cell mRNA sequencing datasets, we saw that 10 of our genes overlapped with some DANs (Figure S5), though the expression overlap varied. *PIG-A* and *Sos* showed only ∼3% overlap, and *Bsf* and *mask* showed 28% and 35% overlap with ^42,43^DANs, respectively.^42,43^ For five of our genes, we were able to examine their expression using available T2A-GAL4 lines.^39–41^ We found variability in the percentage of DANs with overlapping expression, but three of them (*Bsg*, *ClC-c*, and *PIG-A*) overlapped with ∼75% of DANs, *rk* overlapped with ∼45%, and mask overlapped with ∼66% of the examined DAN clusters (Figures S5, S6, 5A, 5B-D, and 5B’-D’). For *clu*, we were able to examine expression using a protein trap line.^44,45^ We saw that clueless protein appears to be enriched in DANs (Figures 5E, 5F-5H and 5F’-5H’).

**Figure 5.**
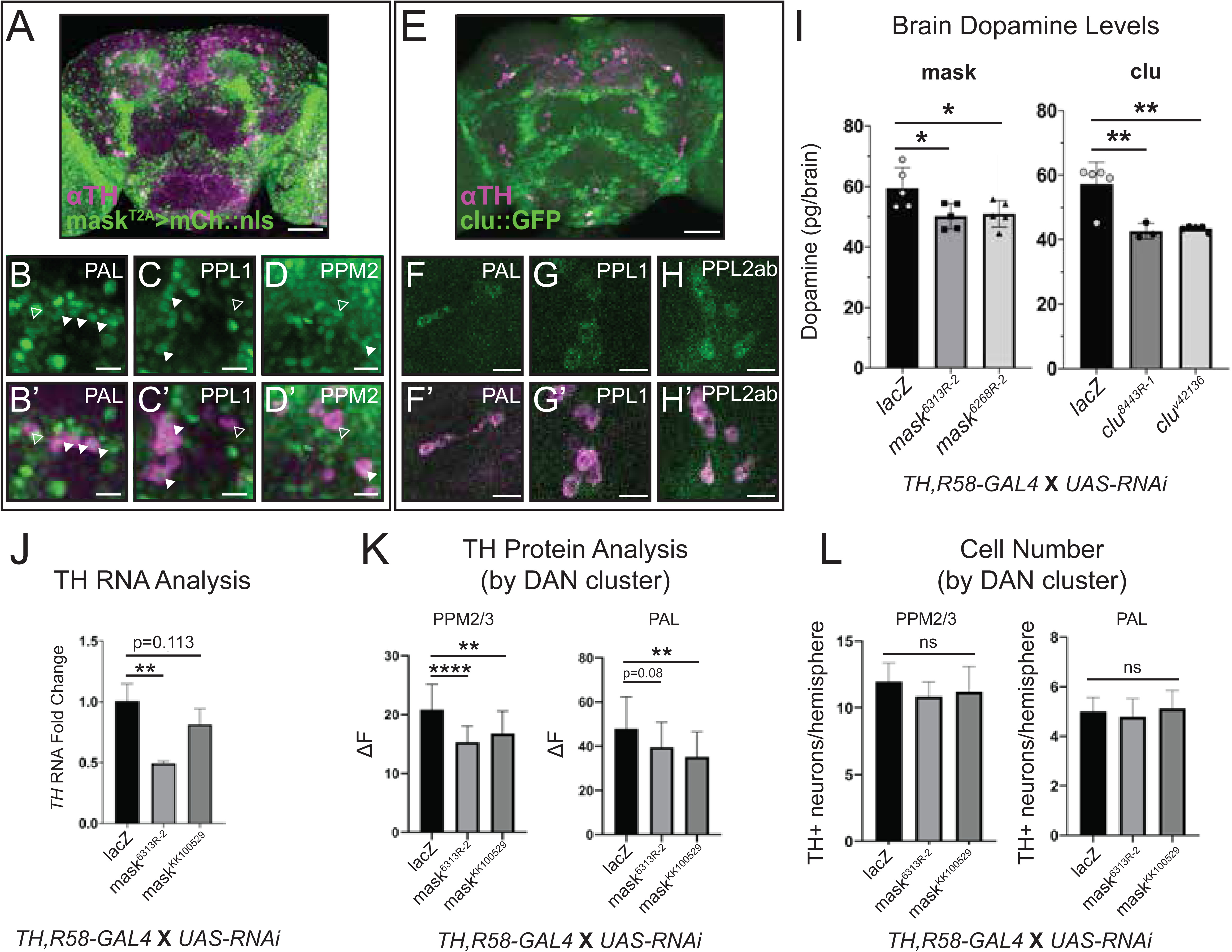
Brain dopamine analysis identifies *mask* and *clu* as novel regulators of dopamine levels in the fly brain. (A) *mask^T2A^>mCh::nls* (represented as green) expression pattern in the adult brain colabelled with a TH antibody staining DANs (magenta), with specific clusters shown (B-D, B’-D’). (E) *clu::GFP* expression pattern in the adult brain colabelled with TH (magenta), with specific clusters shown (F-H, F’-H’). (I) Total dopamine in the adult fly brain upon knockdown of *mask* and *clu*. (J) qRT-PCR of TH upon *mask* knockdown with the recombined DAN (*TH, R58-GAL4*) driver. (K) Quantification of total TH protein level through immunofluorescence in DAN clusters upon knockdown of *mask*. (L) Quantification of TH positive (TH+) neurons in DAN clusters upon knockdown of *mask*. Ordinary one-way ANOVA was done with Dunnett’s multiple comparisons test (I-L). Error bars represent SD. Scale bars are 50 µm in (A) and (E) and 10 µm in all others. Closed arrowheads represent some cells with coexpression between *mask^T2A^>mCh::nls* and TH. Open arrowheads represent cells with TH staining and no *mask^T2A^>mCh::nls* expression.

The *TH-GAL4* line commonly used for dopamine studies misses a large portion of DANs, particularly in the PAM cluster (∼87 DANs missed/∼142 total DANs per hemisphere),^46^ making it not ideal for quantifying total changes in brain dopamine. We recombined the *TH-GAL4* with another driver that hits a large portion of the PAM cluster neurons (*GMR58E02-GAL4 or R58-GAL4)*^18^ generating a recombined line (labelled *TH,R58-GAL4*) that hits ∼95% of DANs (Figure S4). We microdissected adult fly brains and measured dopamine for our 11 gene hits. Most of them showed no change in brain dopamine (Figure S5), but *mask* (*multiple ankyrin repeats single KH domain*) and *clueless* (*clueless*) knockdown reduced total brain dopamine for two independent RNAi lines (^44,45^Figures 5I).

In summary, our fly brain analysis revealed that at least six (*Bsg*, *ClC-c*, *clueless*, *mask, PIG-A*, *rk*) of our gene candidates show co-expression with TH, but only two genes (*clueless* and *mask*) significantly altered brain dopamine upon knockdown in the majority of DANs.

#### *mask* regulates Tyrosine Hydroxylase and alters sleep in a dopamine-dependent manner

To test if *clu* and *mask* knockdown could be due to DAN loss, we quantified the number of DANs in several clusters.^46^ There was no difference in neuron number for any cluster examined (Figures 5L and S6). Since our pigmentation gene study revealed that only the manipulation in TH reduced brain dopamine (Figure 1), we examined if knocking down our genes affect TH. Knockdown of *mask* led to a significant reduction in *TH* mRNA level for one RNAi (50% reduction), with the other RNAi trending (20-30% reduction, Figure 5J). Upon examining TH protein levels in DAN clusters, two clusters showed significant reduction in TH protein level, with other clusters showing a trend in TH protein reduction (PAL and PPM2/3, Figures 5K and S6). To assess behavioral consequences, we performed additional sleep analysis on *mask*. Dopamine is a wake-promoting agent, and complete loss of dopamine in the brain significantly increases sleep.^47,48^ Knocking down *TH* using the *TH-GAL4* did not significantly alter total levels of sleep during light or dark periods (Figure S7), which may be due to the selective expression of the GAL4 or the partial reduction of dopamine (Figure 4C). However, two hours before light onset, there was a significant increase in sleep, accompanied by a reduction in the locomotor activity that anticipates light. Similarly, knockdown of *mask* with the TH-GAL4 showed a highly consistent reduction in light anticipation (Figures 6A, 6B, 6B’, and S8). When we feed the flies L-DOPA, the effect is no longer seen (Figures 6C and S8). Additionally, in *Drosophila* caffeine’s effects on sleep are mediated by dopamine through a point that is upstream of L-DOPA.^49^ Based on this, we suspected that *mask* knockdown would ameliorate the effects of caffeine on sleep, since *mask* appears to reduce the synthesis of *TH*. We saw that caffeine’s effect on total sleep and sleep in the dark is ameliorated upon *mask* knockdown (Figures 6D, 6E, and S8).

**Figure 6.**
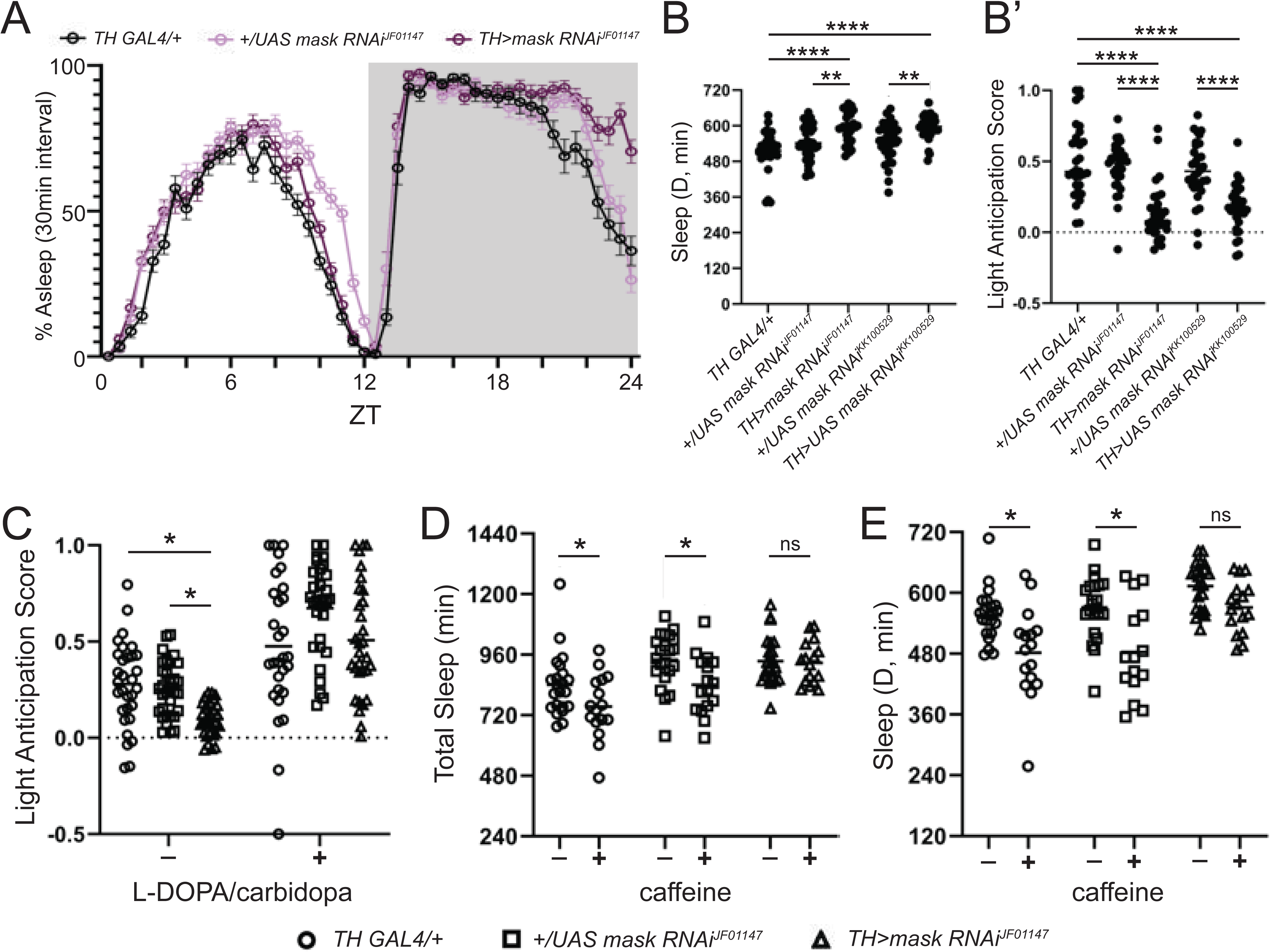
*mask* knockdown reduces TH mRNA and protein and causes sleep phenotypes that align with reduction in TH levels. (A) 24-hr sleep graph (sleep binned in 30-minute intervals) upon mask knockdown using the TH-GAL4. (B) Quantification of total sleep throughout the 12-hr dark period upon mask knockdown (B’) Quantification of light anticipation upon mask knockdown. (C) Light anticipation in mask knockdown flies with L-DOPA feeding. (D-E) Caffeine effects on sleep upon mask knockdown – (D) Total Sleep and (E) Sleep during the 12-hr dark period. For samples with more than one experimental, ordinary one-way ANOVAs were performed with Dunnett’s multiple comparisons. (B, B’, C). For samples with one experimental, individual t-tests were performed (D,E). *=p<0.05, **p<0.01, ****p<0.0001. Error bars represent SD.

Follow up studies on *clueless* showed that *clueless* knockdown increases TH RNA level by about 1.5-2X, with no effect on TH protein level or TH+ neurons (Figure S9), and while there was an increase in sleep across several RNAi lines, there is no light anticipation phenotype. Feeding *TH>clu RNAi* flies L-DOPA was unable to rescue the *clueless* effects on sleep (Figure S9), suggesting that the effects on dopamine and TH may be a secondary effect. Additionally, the antioxidant NACA was also unable to rescue the sleep defects seen upon *clueless* knockdown (Figure S9).

In conclusion, *mask* appears to reduce total dopamine levels and sleep by affecting transcription of *TH*, while *clueless* is acting through an alternative mechanism.

## Discussion

### Validation of 148 novel cuticle pigmentation genes

Our RNAi screened confirmed 153 genes that were previously implicated in *Drosophila* pigmentation – 148 of these genes have not been previously associated with dopamine biology. Since we observed a strong enrichment for genes involved in pigmentation and tyrosine metabolism (Figure S1), our screening method effectively identified genes that are involved in the process of interest.

While prior research has compared genes that overlap with different types of screening strategies (e.g. chemical mutagenesis vs. RNAi), most RNAi phenotypes from large-scale screens have not been validated with alternative RNAi lines or collections. The original source RNAi screen used the lines available at the VDRC stock center.^35,37^ Alternative sources for RNAi lines (i.e. TRiP and NIG) start with different backgrounds, vectors, hairpin lengths, insertion methods, and/or target different areas of the gene.^36,50^ This allows researchers to overcome the off-targeting and weak silencing seen with RNAi analysis, but to our knowledge, there have been no screens or meta-analyses to test consistency from different lines. In this study, we found that ∼46.4% (153/330) of genes with a pigmentation phenotype showed a phenotype in an alternative RNAi, and 87% of those phenotypes showed agreement in pigmentation color, highlighting that pigmentation as a phenotype is relatively consistent (Figure S10).

Interestingly, 85% of our genes had a homolog, whereas the original RNAi screen list (11,619 genes) had 68% of genes with a homology (1.25-fold enrichment, Figure S2). This is striking since dopamine cuticle pigmentation is an invertebrate-specific phenomenon,^12^ and the enrichment suggests this phenotype is relevant for a conserved biological pathway. We propose that cuticle pigmentation may be a “phenolog” ^51^of certain neurological traits in mammals.^51^

### Protein-protein interaction network highlights unexpected gene associations

When we investigated what kind of gene categories were found in the pigmentation screen, we saw clusters that are associated with ubiquitin-proteasome system (UPS), RNA processing, mitochondria, and developmental signaling pathways. Each of these processes could be relevant to dopamine biology. For example, there is some evidence that the UPS system regulates TH levels in the mammals since inhibition of the UPS system has been shown to increase the TH protein in a PC12 rat cell line.^52,53^ In addition, changes in RNA processing could indicate changes in the dopamine synthesis enzymes TH and Ddc since these enzymes undergo RNA processing to create two different isoforms.^54^ The TH isoforms differ in tissue expression and kinetics.^55^ In mammals, post-transcriptional trafficking of TH mRNA and translational regulation has been suggested to be important for its function.^56^

We identified 12 genes that are known to function at the mitochondria, mostly in the electron transport chain (*ND-13B*, *ND-42*, *COX5A*), mitochondrial mRNA translation (*mRpL47*, *mRpL48*, *CG4679*), and mitochondrial protein localization (*CG8728*, *clu*, *mask*, *Mpcp2*). It is well-known that the mitochondria have a key role in dopamine metabolism in mammals since MAO acts in the mitochondria to degrade dopamine. However, there is no known MOA homolog in flies.^12^ The metabolites for oxidation and methylation of dopamine (DOPAC and HVA) have been observed in flies though,^57–60^ and we saw that dopamine degradation enzyme mutants (*ebony*, *tan*, and *speck*)^61,62^ had no effect on dopamine in the brain (Figure 1). It is possible that there is an alternative process that regulates dopamine degradation in the brain, and the mitochondria may be involved.

The last major group of genes identified from our screen was developmental signaling, specifically EGF signaling and Hippo signaling. To date, neither EGF nor Hippo signaling have an established connection with dopamine biology in *Drosophila*. We may have pulled these genes out is because forming the dorso-central thorax is developmentally regulated.^63–65^ However, we did not identify core components of the other key pathways, such as Notch or Hedgehog, suggesting that EGF and Hippo are more specifically involved in this pigmentation process.

### Cuticle pigmentation phenotypes may not reflect changes in dopamine

The field has generally assumed that a darker cuticle reflects an elevated level of dopamine.^15,66^ However, when we measured dopamine from degradation enzyme mutants using HPLC, we saw either no effect on dopamine level or reduced level of dopamine in the head. In addition, when we measured dopamine from the pigmentation genes from the screen, a darker cuticle did not reflect elevated levels of dopamine in the head. While we were able to increase dopamine by 3-5 fold upon TH cDNA overexpression in the head (Figure 4C), this phenomenon was not observable in the brain. One reason could be that there is a selective pressure or a strong feedback mechanism that prevents elevated levels of dopamine in the nervous system. Since oxidized dopamine could be toxic,^67^ this could be a mechanism to prevent cell toxicity. Regardless, changes in cuticle pigmentation do not necessarily reflect changes in dopamine level.

Upon seeing alterations of dopamine in the head for some of the classic pigmentation genes, we tested if this corresponded to changes in the brain. In the case of established dopamine synthesis genes, we saw a mild reduction in dopamine (∼25%) upon knockdown of the rate-limiting enzyme, *TH*, and no effect upon knockdown of *Ddc*. In addition, we saw no effect on brain dopamine from dopamine degradation enzymes. Taken together, this suggests that brain dopamine is under very tight regulation, and effects on the enzymes may be difficult to see in total dopamine level. It also highlights how our approach will likely miss relevant candidates. In addition, upon looking at expression in the brain of dopamine degradation enzymes, we saw they did not overlap well with DANs. It may be that dopamine degradation does not happen in the neurons since it was hypothesized to occur in the glia.^31,68^ However, the expression does not match a broad glial expression either. For the 11 genes from the pigmentation RNAi screen that showed altered levels of dopamine, only 2 (18%) of them showed an effect on dopamine in the brain. These results highlight how head dopamine levels should not be used as a proxy for brain dopamine levels.

### *clu* and *mask* reduce brain dopamine levels through different mechanisms

Once we identified that knockdown of *clueless* and *mask* both reduced dopamine in the brain without affecting the number of DANs, we hypothesized that they may be acting through TH. This is primarily because our initial dopamine metabolism studies showed that only changes in TH significantly affected dopamine levels in the brain.

*mask* encodes a scaffolding molecule that functions in multiple contexts.^69–71^ In this study, *mask* knockdown reduced *TH* mRNA and protein levels, suggesting it acts on TH transcription. Follow-up behavior analysis showed that *mask* knockdown in DANs led to a highly consistent change in sleep. Specifically, *mask* knockdown reduced the number of flies that anticipated light. While fruit flies typically start to wake before light onset,^72^ loss of mask in DANs suppresses this anticipation behavior. Importantly, dopamine has a role in sleep that goes beyond light anticipation, and complete loss of dopamine causes a global increase in sleep in the day and night.^73^ The phenotype we observe is not this severe, which may be due to the GAL4 driver not hitting all DANs or the fact that this knockdown is only reducing dopamine by a modest 15-20% (Figure 5). This is supported by the similar phenotype with the TH RNAi and the DAN driver (TH>TH RNAi, Figure S7). There may be a circadian component as well. It is plausible that dopamine synthesis increases before wake at light onset, and manipulations that affect TH levels may struggle to keep up with the dopamine synthesis required at this time of day.

When we fed *mask* knockdown flies the precursor for dopamine, L-DOPA, this abolished the effect on light anticipation. We suspect that feeding L-DOPA does not override all sleep-dependent effects because clueless sleep phenotypes remained when we fed them L-DOPA (Figure S9). There is evidence that in the fruit fly caffeine affects sleep through TH. Specifically, Nall et al. showed that caffeine’s effect on sleep is absent in TH mutant flies, even if L-DOPA is given.^49^ Since *mask* knockdown affects TH mRNA levels, we tested if mask knockdown would suppress the effects of caffeine. We found that the reduction in total sleep and sleep during the dark period normally seen upon caffeine administration were absent upon mask knockdown. This further supports that mask alters TH levels.

While we do not have any direct evidence for how *mask* may be altering TH levels, *mask* has been implicated in multiple biological pathways that were highlighted from our screen, including EGF signaling (e.g. *Egfr*, *Sev*, Raf), Hippo signaling (e.g. *hpo*, *wts*, *mats*), and mitochondrial biology.^74–77^. Egf and Hippo signaling have well-known transcriptional effects,^78,79^ but whether they may regulate TH transcription directly or indirectly in this context need further investigation. It is also possible that *mask* is acting in several of these pathways at once, which causes it to have a strong additive effect that produces a significant change in dopamine in the whole brain, while knockdown of individual pathways may have more restricted roles (e.g. *Egfr*, *Sos*, *hpo*).

In addition to *mask*, we also saw that knockdown of *clueless* leads to significant reduction in dopamine. However, when we explored the behavioral consequence of *clueless* knockdown, it did not show the same effect on sleep as *mask* or *TH* knockdown. In addition, the sleep phenotype was not rescued by L-DOPA, and molecular biology showed that *clueless* knockdown leads to an increase in TH mRNA with no change in TH protein. Taken together, this indicates that the effect on dopamine from *clueless* knockdown may be a secondary consequence. *clueless* encodes a ribonucleoprotein thought to act at the outer mitochondrial membrane to promote proper formation of protein complexes including mitophagy proteins like Pink1 and Parkin.^80,81^ It also interacts with the ribosome and translocase complex at the mitochondrial membrane suggesting it acts as a regulator for mitochondrial protein translation and import. ^80,82^ Knockdown of *clueless* showed no effect on TH protein level, but since this is a mitochondrial protein, it may be more likely that the effect on dopamine is through the previously proposed roles of mitochondria in regulating dopamine levels.

Importantly, our analysis has focused on changes in dopamine caused by changes in synthesis or degradation. However, dopamine levels can also be regulated by changes dopamine trafficking through regulators like the dopamine transporter (DAT) or the vesicular monoamine transporter (VMAT).^12^ For example, a null allele of the *DAT, fumin*, causes elevated levels of dopamine that lead to behavioral phenotypes, including hyperactivity and reduction in sleep.^48^ However, these mutants do not have any cuticle pigmentation defects. While the role of VMAT in cuticle pigmentation have not been investigated, whether dopamine secretion from the epithelial cells remains a mystery. Therefore, it would be important to further study whether some of the hits from our screen may affect dopamine levels through altering dopamine transport via regulating DAT, VMAT or other mechanisms that regulates dopamine dynamics.

### Some regulators of dopamine levels may act locally

When we examined the expression of some of our pigmentation genes in the brain (Figure S11), we found that many of them do not overlap with all DANs. This indicates that measuring global changes in dopamine may not be best for determining effects on DA for many of these genes. They may require targeted dopamine measurements, perhaps through a fixed or live reporter.^83,84^ It is also possible to assess changes in dopamine using behavior, but this would require detailed assessment for each gene since behaviors are often cluster specific.^85^

In conclusion, our screen of identified novel pigmentation genes, a subset of which were identified as novel regulators of dopamine *in vivo*. Unbiased forward genetic screens in model organisms are powerful ways to identify unanticipated links between distinct biological pathways and provide new molecular handles to study the functional connections between them. Application of such strategies to the regulation of dopamine levels will likely continue to identify novel factors, some of which will impact our understanding of human neurological and neurodevelopmental diseases.

### Limitations of the study

One key limitation to this study is that this screening approach will not capture all regulators of dopamine in the brain. The screen and most of the experiments are based off of RNAi lines. These lines were not verified for the initial and secondary screens. Some RNAis may not work. In addition, the RNAis often do not have the same effect as full mutants. For example, the TH RNAi, which has a known large effect on TH mRNA level (∼90% reduction) still only reduces dopamine by ∼30% in the brain. However, the full mutant for TH removes all dopamine from the brain. In addition to this the initial screen was based on pigmentation, and if a gene that regulates dopamine in the brain is not expressed in the cuticle, it would not have been captured. Our secondary screening uses total dopamine levels in the head and brain. It is likely that mild or cell-type specific changes in dopamine would not be captured through this approach. When we examined *clu* and *mask*, we focused on regulation through the dopamine synthesis enzymes. There are many ways that *clu* and *mask* could regulate dopamine, including through genes such as VMAT and DAT. We cannot rule out dopamine regulation through these alternative mechanisms.

## Supporting information

Supplemental Figures - All

## Resource Availability

All data generated or analyzed during this study are included in this published article and its supplementary information files. This study did not generate any new unique reagents.

## Acknowledgements

We thank the Bloomington *Drosophila* Stock Center (USA), the National Institutes Genetics Fly Stock Center (Japan), *Vienna* Drosophila RNAi Center (Austria) and Dr. Hugo Bellen for providing useful fly stocks and reagents for this project. We would like to thank Drs. Jonathan Andrews, Hugo Bellen, Brigitte Dauwalder, Herman Dierick, Linsday Goodman, Oguz Kanca, Kartik Venkatachalam, Michael Wangler, Sheng Zhang, Huda Zoghbi, and the Sehgal lab for useful suggestions, discussions and advice on the project. This work was supported by startup funds to Dr. Shinya Yamamoto from the Jan and Dan Duncan Neurological Research Institute at Texas Children’s Hospital and the Department of Molecular and Human Genetics at Baylor College of Medicine, by the IRACDA program at the University of Pennsylvania (K12GM081259) for Dr. Samantha Deal, and by Howard Huges Medical Institute (HHMI) for Dr. Amita Sehgal. Confocal microscopy at BCM was supported in part by the Intellectual and Developmental Disabilities Research Center (IDDRC, U54HD083092) from the Eunice Kennedy Shriver National Institute of Child Health & Human Development.

## Authors Contributions

S.Y., E.S.S., A.S., and S.L.D. conceived the experiments. S.Y. and S.L.D. wrote the manuscript. S.Y., E.S.S., D.B., and S.L.D. conducted the pigmentation screen and DAM behavior analysis screen. S.L.D. and K.W. conducted the pigmentation mutant analysis experiments. S.L.D., S.B.G, H.D-S, Y.F., and K.W. conduced the HPLC analysis. S.L.D. and Y.F. performed confocal imaging. S.L.D. conducted all data analysis. All authors participated in the critical analysis of the manuscript.

## STAR★Methods

### Experimental Model and Study Participant Details

#### Fly Maintenance and Stocks

Flies (*Drosophila melanogaster*) were reared at 25°C on a 12:12 Light/Dark (LD) and fed a molasses-based food source unless otherwise specific. Different temperatures were used to achieve different levels of gene knockdown or overexpression, which is documented for each experiment. Flies were transferred to a new vial 0-2 days post-eclosion (dpe) for age-appropriate analysis. Experiments were run on 3-7 dpe flies unless otherwise stated. Mutant and transgenic trains were obtained from Bloomington *Drosophila* Stock Center (BDSC, https://bdsc.indiana.edu/), Vienna Drosophila Research Center (VDRC, https://www.viennabiocenter.org/vbcf/vienna-drosophila-resource-center/), the Japanese National Institute of Genetics (https://shigen.nig.ac.jp/fly/nigfly/), or were gifts from scientists in the field or generated in house (S1 Table – S5 Table).

#### Fly Crosses

For the cuticle screen, *pnr-GAL4*^86^ was crossed to the specified UAS-RNAi line (see S1 Table) at 29°C, 25°C, and/or 18°C and examined after pigmentation is completed (>1 dpe). For the behavior and HPLC analysis *TH-GAL4* females (also called *ple-GAL4*, BDSC stock #8848) were crossed to the specified RNAi line (see S2 Table). For all HPLC analysis, the controls were specific to the tested UAS-RNAi line.^87^ Thus, the TRiP collection of RNAi lines are compared to the *UAS-mCherry(mCh) RNAi* line (BDSC #35785) and the NIG collection of RNAi lines as well as the VDRC RNAi lines are compared to the *UAS-lacZ RNAi* line. For the HPLC analysis performed on brains, we recombined the *TH-GAL4* with another DAN driver *GMR58E02-GAL4* (BDSC #41347), which expresses in a large group DANs in the PAM cluster.^27^ These flies are referred to as *TH,R58-GAL4* throughout the article. The *TH,R58-GAL4* line was also used for qPCR and TH protein analysis via immunohistochemistry. For gene expression analysis *UAS-mCh::nls* (BDSC #38424) females were collected and crossed with respective T2A-GAL4 lines (see S5 Table). Female and males were both imaged at ∼5-7dpe. For the follow-up sleep studies, the *UAS-RNAi* males were crossed with *TH-GAL4* virgins for the experimental. For the controls *w^11^*^18^ females were crossed to males from either the *TH-GAL4* and *UAS-RNAi* lines independently.

### Method Details

#### Notum Dissection and Imaging

For thorax analysis and dissection, males and females were both selected, though differences were not generally observed, and females are shown here. Flies that were 3-5 dpe were collected and placed in 70% EtOH. Notum dissection and imaging was previously published.^88^ In summary, the thorax was dissected by removing the legs, abdomen, and head. Then the thorax was cut such that a hole was on the ventral side to allow solution to pass into the thorax. These dissected thoraces were placed in 10% KOH for 10 minutes at 90°C on a heat block. Then, the KOH was removed and replaced with 70% EtOH solution. They were mounted on a slide prepared with tape on two sides in mounting media (50% glycerol, 50% EtOH). The thoraces were imaged on a stereo microscope (Leica MZ16) using OPTRONICS® MicroFIRE camera. The images are z-stack brightfield images taken and collapsed using extended depth of field in Image-Pro Plus 7.0 and In-Focus (Version 1.6).

#### Brain Expression Analysis

Brain dissections and imaging were performed as previously reported.^89^ In summary, the brains were dissected at 3-7 dpe in ice cold PBS and then fixed in 4% PFA in 0.5% PBST for 20 minutes. Then, they were washed with a quick wash in 0.5% PBST followed by three 10-20-minute washes in 0.5% PBST while on a rotator. The samples were placed in primary antibody solution [anti-Tyrosine Hydroxylase 1:500, PelFreez Biologicals, rabbit, P40101; anti-elav 1:100, Developmental Studies Hybridoma Bank, rat, 7E8A10 in solution (5% Normal Donkey Serum, 0.1% NaN_3_ in 0.5% PBST)] at 4°C for 3 days. Then, the samples were washed with a quick wash in 0.5% PBST followed by three 10-20-minute washes in 0.5% PBST while on a rotator. After the last wash, the samples were placed in a secondary antibody solution [anti-Rabbit 1:200 (Thermo Fisher Sci., Alexa-647, A-21208); anti-rat 1:200 (Thermo Fisher Sci., Alexa-488, A-27040) in solution (5% Normal Donkey Serum, 0.1% NaN_3_ in 0.5% PBST)] for two hours at room temperature. The samples were washed again with one quick wash followed by three 10-20 minutes washes in 0.5% PBST on a rotator. Then, the brains were mounted in Vectashield mounting media and imaged on a confocal microscope (Zeiss LSM 710 or Zeiss LSM 880). All images shown in this manuscript are Z-projection images that were generated using the ZEN software (Zeiss). Quantification of co-expression with DANs and the T2A-GAL4 lines was done by counting how many individual TH+ neurons for each cluster had a cell body labeled by the *T2A-GAL4>mCherry::nls* signal. This quantification was done per hemisphere. The single cell sequencing expression analysis was done based on the DAN cluster from adult fruit fly data found in the Fly Cell Atlas, specifically using the data from Davie, Jannssens and Koldere et al., 2018.^91^

#### RNAi-based Pigmentation Screen

We identified pigmentation gene hits from a primary screen performed in Jurgen Knoblich’s lab using the Vienna Drosophila Resource Center collection of RNAi lines by accessing their public RNAi screen database (https://bristlescreen.imba.oeaw.ac.at/start.php). The original screen was performed on 20,262 RNAi lines, encompassing 11,619 genes (∼82% of protein-coding genes).^22^ We selected genes that showed a cuticle pigmentation score for any RNAi line tested (gene scores ranged from two to ten). To validate pigmentation defects, we selected genes that had RNAi lines within the Fly National Institute of Genetics (NIG, 220 genes, 426 RNAi lines) or the Harvard Transgenic RNAi Project (TRiP, 221 genes, 292 RNAi lines) collections. Each RNAi line was tested at 29°C and 25°C. If there was no phenotype no further testing was performed. If it was lethal, we tested it at 18°C.

#### Classification of Cuticle Phenotypes

Phenotypes were scored using a qualitative scoring system. This system used *UAS-TH RNAi* and *UAS-ebony RNAi*, as a baseline for “strong”. Then, lines were placed along the spectrum as mild, moderate, or strong. Phenotypes were scored by two independent observers and differences were settled by a third independent observer. If there was variability with one line or if there was variability amongst lines, this might appear as mild-moderate, moderate-strong, or mild-strong. In these cases, the strongest phenotype observed was documented as the recorded phenotype for a given gene.

#### Protein-protein interaction network and gene ontology analysis

The protein-protein interaction network was generated using STRING (search tool for recurring instances of neighboring genes, https://string-db.org/). For our data set we included these sources of interactions: gene neighborhoods (genes that are found close together across species),^90^ curated databases (publicly available databases of protein interactions), and experimental evidence (known complexes and pathways from curated sources).^92,93^ Any genes that did not interact with other genes from our screen were not included in the interaction network. This database was last accessed on 05/05/2023. The lines between genes represents the confidence in the interactions.

The gene ontology analysis was generated using GOrilla (Gene Ontology enRIchment anaLysis and visuaLizAtion tool, https://cbl-gorilla.cs.technion.ac.il/). The 153 pigmentation genes hits were included as target genes, and the background genes were the entire VDRC RNAi collection included in the Mummery-Widmer et al. 2009 screen. Only the GO terms with a p-value greater than 0.001 and a fold enrichment of greater than three were included in the analysis. Redundant terms were removed using Revigo (http://revigo.irb.hr/) with a stringency of 0.7. Those that have a fold enrichment greater than five were included in S1 Figure.

#### Human Neurological Disease Gene Classification

Homologs for each fly gene were identified using DIOPT (DRSC Integrative Ortholog Prediction Tool, v8.0, https://www.flyrnai.org/diopt).^94^ Genes were classified as orthologs if they had a DIOPT score≥3. The human genes were scored as neurological disease-causing genes if they had an OMIM (Online Mendelian Inheritance of Man, last accessed 01/01/2023, https://www.omim.org/) disease association with any documented neurological phenotype.^95^ Genes were also classified as human neurological disease genes if they were in the Simons Simplex Collection of Autism Spectrum Disorder gene list (last accessed 06/01/2022, https://gene.sfari.org/) and had a score equal to or less than 3 (1, 2, or 3) and/or syndromic (S).^96^

#### Behavior Analysis using the Drosophila Activity Monitor

##### Behavior screen

For the behavior screen, the crosses were set in a 12:12 LD chamber at 25°C and transferred every 2-3 days to increase the number of progenies. Upon eclosion, flies were transferred to 29°C and kept for 2-3 more days before testing. Individual tubes (PPT5×65 Polycarbonate, Trikinetics Inc, USA) appropriate for use with the *Drosophila* Activity Monitor (DAM, Model DAM2 for 5mm tubes, TriKinetics Inc, USA) were loaded with approximately ½” worth of molasses-based food. The end of the tube with food was sealed by placing the vial into Paraplast® (Sigma-Aldrich) wax three times and allowing it to dry. Once they were dry, individual male flies were placed into each vial, the vial was placed into the DAM recording chamber, and it was sealed using a 5mm tube cap (CAP5, TriKinetics Inc, USA) with a hole stuffed with cotton. For each genotype, 16 individual males were run, except in a few cases where we were unable to get that many flies. In which case, as many living males as possible were run. The flies were loaded into a 12:12 LD chamber at 29°C and monitored for 3 days. Then, the flies were removed and dead flies were eliminated from analysis.

Locomotion and sleep analysis was run on a 24-hour period, which started at the first onset of light after the flies were placed in the chamber (i.e. 18-24 hours after being placed in the chamber). Locomotion was calculated based on the number of beam crossings over the full 24 hours, the 12 hours of light, and the 12 hours of dark. Sleep was classified as 5 minutes without any beam crossings. Total Sleep was quantified as total time spent sleeping over the 12 hours of light and the 12 hours of dark. Sleep latency was quantified as the amount of time after light onset or dark onset before a 5-minute period of sleep. Sleep Bout Length was quantified as the average length of sleep bouts (>5-minute period of sleep) during the 12 hours of light and 12 hours of dark.

“Outliers” were classified as lines that were more than two standard deviations from the mean of all lines for any of the phenotypes scored (24-hour locomotion, 12-hour light locomotion, 12-hour dark locomotion, 12-hour light total sleep, 12-hour dark total sleep, 12-hour light sleep bout length, 12-hour dark sleep bout length, sleep latency during the light, or sleep latency during the dark).

##### Mask sleep study

For these studies, similar tools and settings were used with these exceptions. Upon eclosion flies were transferred and kept at 25°C in group housing. For the behavior without drugs, individual males were placed in tubes with sucrose food (2% agar + 5% sucrose) at 3-6 dpe and behavioral analysis was collected and analyzed for the following five days. Sleep was then averaged across the five days for each individual fly before statistical analysis was performed. For the behavior with drugs, similar conditions were followed, except data was collected on the 2-6 days after they were loaded into behavior tubes to allow the drug to take effect. The following drug doses were used for their respective experiments (L-DOPA: 3mg/mL L-DOPA + 12.5 µg/mL carbidopa, caffeine: 0.5 mg/mL caffeine, NACA: 40 µg/mL N-acetylcysteine amide antioxidant). Drugs were dissolved directly into the sucrose food except for carbidopa which was dissolve in H_2_O at 1:50x concentration and then added into the sucrose food.

#### High Performance Liquid Chromatography (HPLC) analysis

##### HPLC System

The HPLC used is an Antec® Scientific product with a LC110S pump, SYSTEC OEM MINI Vacuum Degasser, AS110 autosampler, a SenCell flow cell with salt bridge reference electrode in the Decade Lite. Data was collected and processed using DataApex Clarity™ chromatography software. The column used is chosen to work well for neurotransmitters (Acquity UPLC BEH C18 Column, 130Å, 1.7 µm, 1 mm X 100 mm with Acquity In-Line 0.2 µm Filter). The mobile phase was a 6% Acetonitrile mobile phase optimized for our samples (74.4 mg NA_2_EDTA·2H_2_0, 13.72 mL 85% w/v phosphoric acid, 42.04 g citric acid, 1.2 g OSA, 120 mL acetonitrile, H_2_0 up to 2L, pH=6.0 using 50% NaOH solution). The mobile phase was degassed for 10 minutes using the Bransonic® Ultrasonic Bath before being loaded into the HPLC machine.

The standards for HPLC were made by generating master stocks of 100 mM dopamine and 10 mM serotonin diluted in MilliQ H_2_O, which were kept at 4°C. The day of HPLC analysis, the master stocks were diluted to produce standards ranging from 5-100 nM for 5-HT and 5-1000 nM for dopamine. Sample concentrations of dopamine (Sigma-Aldrich, Cat#H8502) and serotonin (Sigma-Aldrich, Cat#H7752) were calculated based on standards run in the same batch. Standards were compared to standards run on other days to assess for overall performance.

##### Sample Preparation

For the HPLC on fly heads and brain, we adapted and modified the protocol previously published.^97,98^ Crosses were reared at 29°C and flies were transferred into new tubes 0-2 days post eclosion (dpe). They remained at 29°C until they were 3-5 dpe. Female heads were collected by anesthetizing flies with CO_2_ and cutting their heads off with a razor blade. Samples were collected between ZT04-ZT08. The heads were placed into 60 µL of 50 mM citrate acetate (pH=4.5) and either used for analysis that day or frozen at −20°C. Five heads were used per sample, and ∼10 samples were run per genotype. The day of HPLC analysis, the samples were thawed and grounded using a pestle for 30 seconds (Cordless Pestle Motor and Fisherbrand™ Disposable Pellet Pestle for 1.5mL tube). Then the samples were spun down at 13,000 rpm for 10 minutes. The supernatant was removed and placed into a new vial. 10 µL from each sample was used for the Bradford Protein Analysis Assay. The rest of the sample solution (∼40-50 µL) was loaded into a vial (300 µL Polypropylene Sample Vials with 8mm Snap Caps) for HPLC analysis.

For HPLC analysis on fly brains, flies were collected 3-7 dpe. Then, 10 fly brains (5 male, 5 female) were dissected in ice cold PBS. Right after dissection, the brains were transferred to 60uL of ice cold 50mM citrate acetate (pH=4.5). Samples were frozen at −20°C and ran within 4 weeks of collection. The day of HPLC analysis, the samples were homogenized similarly to the fly heads, though for the Protein Assay, 20 µL of sample was used. Then, the remaining solution was loaded for HPLC analysis.

##### Bradford Protein Analysis Assay

The Bio-Rad Bradford Assay was used for colorimetric scoring of total protein. See product details for full description (Bio-Rad Protein Assay Kit I #5000001). Briefly, the dye reagent was diluted 1:4 in MilliQ H_2_O. Then, the solution was filtered through a 0.22 µm SFCA Nalgene filter. 10 µL of each protein sample (standard or fly sample) was placed into single wells of a 96-well plate. Then, 200 µL of diluted 1:4 dye reagent was added to each well. The samples rested for 30-45 minutes and then absorbance was measured using BMG Labtech FLUOstar OPTIMA microplate reader. Protein measurements for each sample were calculated based on standards ran on the same plate.

#### RNA expression analysis

Flies were raised at 29°C until 5-7 dpe, then heads were collected by cutting them off with a razor. Twenty heads were collected per sample with a mixture of males and females and placed directly on dry ice. Samples were kept on dry ice or at −80°C until RNA isolation. For RNA isolation samples were homogenized in 200μL of TRIzol®. Then 800μL of TRIzol® was added and the samples were mixed by pipetting. The samples incubated for 5 minutes at room temperature and were then spun down at 12,000 rpm for 10 minutes at 4°C. The supernatant was removed and 200μL of chloroform was added. The samples were mixed by shaking and then incubated at room temperature for 3 minutes. Then, they were spun at 10,000 rpm for 15 minutes at 4°C, and the aqueous layer was collected and placed in a new tube. 500μL of isopropanol was added and the samples were left to incubate for 10 minutes at room temperature. The samples were spun at 10,000 rpm for 10 minutes at 4°C and the supernatant was removed. The pellet was washed with 1mL of 75% EtOH and centrifuged at 5,000 rpm for 5 minutes. All EtOH was removed and the pellet dried at room temperature for 10 minutes. Then, the pellet was dissolved in 100μL of H_2_O. RNA content and quality was measured using DeNovix DS-11 Fx spectrophotometer/fluorometer. cDNA reverse transcription was performed according to the iScript™ reverse transcriptase Bio-Rad kit and 2 µL of RNA sample was added in a 10 µL reaction for both the reverse and no reverse transcriptase reaction. Samples were measured for content and quality and then they were diluted to 100ng/μL. qPCR was performed using Bio-Rad iQ SYBR Green Supermix and measured on Bio-Rad CFX96™ Real-Time System. The TH primers used Forward: 5’-ATGTTCGCCATCAAGAAATCCT-3’ and Reverse: 5’-GGGTCTCGAAACGGGCATC-3’, and the control primers were for Rpl32 were Forward: 5’-ATGCTAAGCTGTCGCACAAATG-3’ and Reverse: 5’-GTTCGATCCGTAACCGATGT-3’. For each sample, two technical replicates were run for the reverse transcriptase reaction, and the final quantity was based on the average of the two. If the no reverse transcriptase reaction produced a product that was a Cq within ten, the sample was discarded.

#### DAN and TH Quantification

Samples were stained and imaged like the brain expression analysis, except the primarily antibody solution included an Elav antibody [anti-Tyrosine Hydroxylase 1:500, PelFreez Biologicals, rabbit, P40101; anti-Elav 1:100, Developmental Studies Hybridoma Bank, rat, 7E8A10 in solution (5% Normal Donkey Serum, 0.1% NaN_3_ in 0.5% PBST)]. Confocal imaging was performed as described above. Once the samples were imaged, anterior, posterior, and whole brain projections were generated that included all fluorescent signals from the anterior (PAL and PAM clusters) and posterior (PPL1, PPM2, PPM3, PPL2ab) neuron clusters of interest. The number of neurons per cluster (excluding the PAM cluster) were counted for each sample. PPM2 and PPM3 clusters were counted together. For TH protein quantification samples were assessed using ImageJ. A boundary was drawn around each individual cluster and quantified. Then, the local background for that cluster was subtracted from the cluster to give the fluorescence for the given sample. Each hemisphere was treated as a separate sample.

### Quantification and Statistical Analysis

Statistical analysis was performed using Graphpad Prism 10. Unless otherwise stated, all data was subjected to a ROUT outlier test, where all outliers were removed, and then a one-way ANOVA was performed. Each of the experimental samples were compared to the control for the given samples. In the cases where there were only two samples (i.e. control and experimental), a t-test was performed.

For the HPLC screen on heads, the samples were normalized to the controls run on that day. The TRiP collection was normalized to *UAS-mCh RNAi* control and the NIG collection was normalized to the *UAS-lacZ RNAi* control.

## Supplemental Information

Document S1. Figures S1-S11

**Table S1. Screen results by RNAi line: Pigmentation, Drosophila Activity Monitor, HPLC, and Prioritization, related to Figures 2 - 4.**

**Table S2. Experimental models: Organisms/strains, related to Key resources table**

---

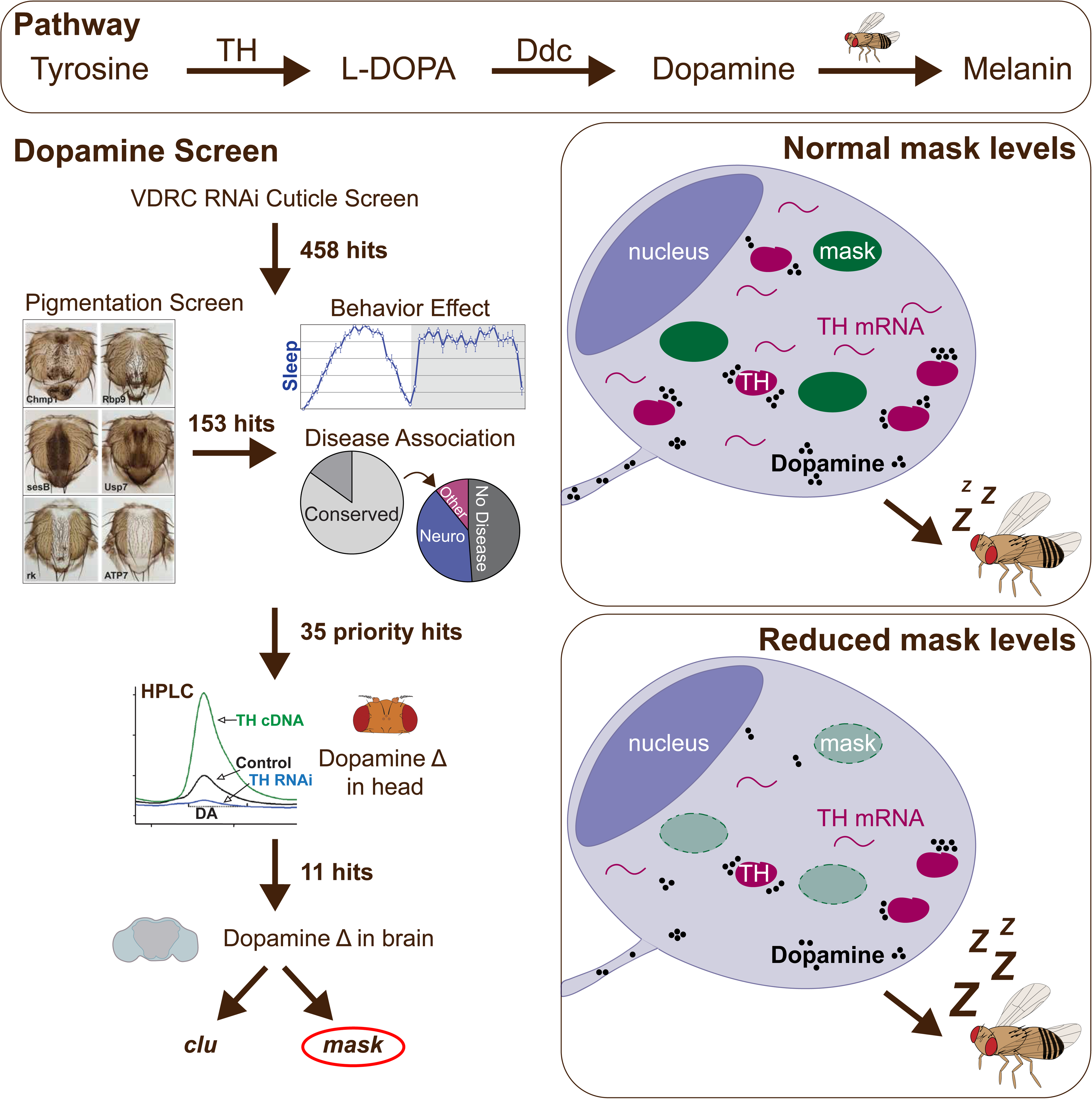

## References

1. Radwan, B., Liu, H., and Chaudhury, D. (2019). The role of dopamine in mood disorders and the associated changes in circadian rhythms and sleep-wake cycle. Brain Res 1713, 42–51. 10.1016/J.BRAINRES.2018.11.031.

2. Grace, A.A. (2016). Dysregulation of the dopamine system in the pathophysiology of schizophrenia and depression. Nat Rev Neurosci 17, 524–532. 10.1038/NRN.2016.57.

3. Nutt, D.J., Lingford-Hughes, A., Erritzoe, D., and Stokes, P.R.A. (2015). The dopamine theory of addiction: 40 years of highs and lows. Nat Rev Neurosci 16, 305–312. 10.1038/NRN3939.

4. Church, F.C. (2021). Treatment Options for Motor and Non-Motor Symptoms of Parkinson’s Disease. Biomolecules 11. 10.3390/BIOM11040612.

5. Armstrong, M.J., and Okun, M.S. (2020). Diagnosis and Treatment of Parkinson Disease: A Review. JAMA 323, 548–560. 10.1001/JAMA.2019.22360.

6. López-Cruz, L., Miguel, N.S., Carratalá-Ros, C., Monferrer, L., Salamone, J.D., and Correa, M. (2018). Dopamine depletion shifts behavior from activity based reinforcers to more sedentary ones and adenosine receptor antagonism reverses that shift: Relation to ventral striatum DARPP32 phosphorylation patterns. Neuropharmacology 138, 349–359. 10.1016/J.NEUROPHARM.2018.01.034.

7. Steiner, H., and Kitai, S.T. (2001). Unilateral striatal dopamine depletion: time-dependent effects on cortical function and behavioural correlates. Eur J Neurosci 14, 1390–1404. 10.1046/J.0953-816X.2001.01756.X.

8. Rohwedder, A., Wenz, N.L., Stehle, B., Huser, A., Yamagata, N., Zlatic, M., Truman, J.W., Tanimoto, H., Saumweber, T., Gerber, B., et al. (2016). Four Individually Identified Paired Dopamine Neurons Signal Reward in Larval Drosophila. Curr Biol 26, 661–669. 10.1016/J.CUB.2016.01.012.

9. Budnik, V., and White, K. (1987). Genetic dissection of dopamine and serotonin synthesis in the nervous system of Drosophila melanogaster. J Neurogenet 4, 309–314. 10.3109/01677068709167191.

10. Livingstone, M.S., and Tempel, B.L. (1983). Genetic dissection of monoamine neurotransmitter synthesis in Drosophila. Nature 303, 67–70. 10.1038/303067A0.

11. Meiser, J., Weindl, D., and Hiller, K. (2013). Complexity of dopamine metabolism. Cell Commun Signal 11. 10.1186/1478-811X-11-34.

12. Yamamoto, S., and Seto, E.S. (2014). Dopamine dynamics and signaling in Drosophila: an overview of genes, drugs and behavioral paradigms. Exp Anim 63, 107–119. 10.1538/EXPANIM.63.107.

13. Wittkopp, P.J., True, J.R., and Carroll, S.B. (2002). Reciprocal functions of the Drosophila yellow and ebony proteins in the development and evolution of pigment patterns. Development 129, 1849–1858. 10.1242/DEV.129.8.1849.

14. True, J.R., Edwards, K.A., Yamamoto, D., and Carroll, S.B. (1999). Drosophila wing melanin patterns form by vein-dependent elaboration of enzymatic prepatterns. Curr Biol 9, 1382–1391. 10.1016/S0960-9822(00)80083-4.

15. Spana, E.P., Abrams, A.B., Ellis, K.T., Klein, J.C., Ruderman, B.T., Shi, A.H., Zhu, D., Stewart, A., and May, S. (2020). speck, First Identified in Drosophila melanogaster in 1910, Is Encoded by the Arylalkalamine N-Acetyltransferase (AANAT1) Gene. G3 (Bethesda) 10, 3387–3398. 10.1534/G3.120.401470.

16. Gervasi, N., Tchénio, P., and Preat, T. (2010). PKA dynamics in a Drosophila learning center: coincidence detection by rutabaga adenylyl cyclase and spatial regulation by dunce phosphodiesterase. Neuron 65, 516–529. 10.1016/J.NEURON.2010.01.014.

17. Heisenberg, M., Borst, A., Wagner, S., and Byers, D. (1985). Drosophila mushroom body mutants are deficient in olfactory learning. J Neurogenet 2, 1–30. 10.3109/01677068509100140.

18. Jenett, A., Rubin, G.M., Ngo, T.T.B., Shepherd, D., Murphy, C., Dionne, H., Pfeiffer, B.D., Cavallaro, A., Hall, D., Jeter, J., et al. (2012). A GAL4-driver line resource for Drosophila neurobiology. Cell Rep 2, 991–1001. 10.1016/J.CELREP.2012.09.011.

19. Frighetto, G., Zordan, M.A., Castiello, U., Megighian, A., and Martin, J.R. (2022). Dopamine Modulation of Drosophila Ellipsoid Body Neurons, a Nod to the Mammalian Basal Ganglia. Front Physiol 13. 10.3389/FPHYS.2022.849142.

20. Xie, T., Ho, M.C.W., Liu, Q., Horiuchi, W., Lin, C.C., Task, D., Luan, H., White, B.H., Potter, C.J., and Wu, M.N. (2018). A Genetic Toolkit for Dissecting Dopamine Circuit Function in Drosophila. Cell Rep 23, 652–665. 10.1016/j.celrep.2018.03.068.

21. Clark, I.E., Dodson, M.W., Jiang, C., Cao, J.H., Huh, J.R., Seol, J.H., Yoo, S.J., Hay, B.A., and Guo, M. (2006). Drosophila pink1 is required for mitochondrial function and interacts genetically with parkin. Nature 441, 1162–1166. 10.1038/NATURE04779.

22. Mummery-Widmer, J.L., Yamazaki, M., Stoeger, T., Novatchkova, M., Bhalerao, S., Chen, D., Dietzl, G., Dickson, B.J., and Knoblich, J.A. (2009). Genome-wide analysis of Notch signalling in Drosophila by transgenic RNAi. Nature 458, 987–992. 10.1038/NATURE07936.

23. Pendleton, R.G., Rasheed, A., Sardina, T., Tully, T., and Hillman, R. (2002). Effects of Tyrosine Hydroxylase Mutants on Locomotor Activity in Drosophila: A Study in Functional Genomics.

24. Wright, T.R.F., Bewleyz, G.C., and Sherald3, A.F. (1976). THE GENETICS OF DOPA DECARBOXYLASE IN DROSOPHILA MELANOGASTER. 11. ISOLATION AND CHARACTERIZATION OF DOPA-DECARBOXY LASE-DEFICIENT MUTANTS AND THEIR RELATIONSHIP TO THE a-METHYL-DOPA-HYPERSENSITIVE MUTANTS.

25. Ni, J.Q., Liu, L.P., Binari, R., Hardy, R., Shim, H.S., Cavallaro, A., Booker, M., Pfeiffer, B.D., Markstein, M., Wang, H., et al. (2009). A Drosophila resource of transgenic RNAi lines for neurogenetics. Genetics 182, 1089–1100. 10.1534/GENETICS.109.103630.

26. Pan, Y., Li, W., Deng, Z., Sun, Y., Ma, X., Liang, R., Guo, X., Sun, Y., Li, W., Jiao, R., et al. (2022). Myc suppresses male-male courtship in Drosophila. EMBO J 41. 10.15252/EMBJ.2021109905.

27. Riemensperger, T., Issa, A.R., Pech, U., Coulom, H., Nguyễn, M.V., Cassar, M., Jacquet, M., Fiala, A., and Birman, S. (2013). A single dopamine pathway underlies progressive locomotor deficits in a Drosophila model of Parkinson disease. Cell Rep 5, 952–960. 10.1016/J.CELREP.2013.10.032.

28. Brand, H., and Perrimon, N. (1993). Targeted gene expression as a means of altering cell fates and generating dominant phenotypes.

29. Bayersdorfer, F., Voigt, A., Schneuwly, S., and Botella, J.A. (2010). Dopamine-dependent neurodegeneration in Drosophila models of familial and sporadic Parkinson’s disease. Neurobiol Dis 40, 113–119. 10.1016/J.NBD.2010.02.012.

30. Hovemann, B.T., Ryseck, R.P., Walldorf, U., Störtkuhl, K.F., Dietzel, I.D., and Dessen, E. (1998). The Drosophila ebony gene is closely related to microbial peptide synthetases and shows specific cuticle and nervous system expression. Gene 221, 1–9. 10.1016/S0378-1119(98)00440-5.

31. Richardt, A., Rybak, J., Störtkuhl, K.F., Meinertzhagen, I.A., and Hovemann, B.T. (2002). Ebony protein in the Drosophila nervous system: optic neuropile expression in glial cells. J Comp Neurol 452, 93–102. 10.1002/CNE.10360.

32. Yamamoto-Hino, M., Yoshida, H., Ichimiya, T., Sakamura, S., Maeda, M., Kimura, Y., Sasaki, N., Aoki-Kinoshita, K.F., Kinoshita-Toyoda, A., Toyoda, H., et al. (2015). Phenotype-based clustering of glycosylation-related genes by RNAi-mediated gene silencing. Genes to Cells 20, 521–542. 10.1111/gtc.12246.

33. Riemensperger, T., Isabel, G., Coulom, H., Neuser, K., Seugnet, L., Kume, K., Iché-Torres, M., Cassar, M., Strauss, R., Preat, T., et al. (2011). Behavioral consequences of dopamine deficiency in the Drosophila central nervous system. Proc Natl Acad Sci U S A 108, 834–839. 10.1073/pnas.1010930108.

34. Chiu, J.C., Low, K.H., Pike, D.H., Yildirim, E., and Edery, I. (2010). Assaying locomotor activity to study circadian rhythms and sleep parameters in Drosophila. Journal of Visualized Experiments. 10.3791/2157.

35. Dietzl, G., Chen, D., Schnorrer, F., Su, K.C., Barinova, Y., Fellner, M., Gasser, B., Kinsey, K., Oppel, S., Scheiblauer, S., et al. (2007). A genome-wide transgenic RNAi library for conditional gene inactivation in Drosophila. Nature 448, 151–156. 10.1038/NATURE05954.

36. Ni, J.Q., Zhou, R., Czech, B., Liu, L.P., Holderbaum, L., Yang-Zhou, D., Shim, H.S., Tao, R., Handler, D., Karpowicz, P., et al. (2011). A genome-scale shRNA resource for transgenic RNAi in Drosophila. Nat Methods 8, 405–407. 10.1038/NMETH.1592.

37. Vissers, J.H.A., Manning, S.A., Kulkarni, A., and Harvey, K.F. (2016). A Drosophila RNAi library modulates Hippo pathway-dependent tissue growth. Nature Communications 2016 7:1 *7*, 1–6. 10.1038/ncomms10368.

38. Shafer, O.T., and Keene, A.C. (2021). The Regulation of Drosophila Sleep. Preprint at Cell Press, 10.1016/j.cub.2020.10.082 10.1016/j.cub.2020.10.082.

39. Diao, F., Ironfield, H., Luan, H., Diao, F., Shropshire, W.C., Ewer, J., Marr, E., Potter, C.J., Landgraf, M., and White, B.H. (2015). Plug-and-play genetic access to drosophila cell types using exchangeable exon cassettes. Cell Rep 10, 1410–1421. 10.1016/J.CELREP.2015.01.059.

40. Lee, P.T., Zirin, J., Kanca, O., Lin, W.W., Schulze, K.L., Li-Kroeger, D., Tao, R., Devereaux, C., Hu, Y., Chung, V., et al. (2018). A gene-specific T2A-GAL4 library for Drosophila. Elife 7. 10.7554/ELIFE.35574.

41. Kanca, O., Zirin, J., Garcia-Marques, J., Knight, S.M., Yang-Zhou, D., Amador, G., Chung, H., Zuo, Z., Ma, L., He, Y., et al. (2019). An efficient CRISPR-based strategy to insert small and large fragments of DNA using short homology arms. Elife 8, e51539. 10.7554/eLife.51539.

42. Janssens, J., Aibar, S., Taskiran, I.I., Ismail, J.N., Gomez, A.E., Aughey, G., Spanier, K.I., Rop, F.V. De, González-Blas, C.B., Dionne, M., et al. (2022). Decoding gene regulation in the fly brain. Nature 601, 630–636. 10.1038/S41586-021-04262-Z.

43. Li, H., Janssens, J., de Waegeneer, M., Kolluru, S.S., Davie, K., Gardeux, V., Saelens, W., David, F.P.A., Brbić, M., Spanier, K., et al. (2022). Fly Cell Atlas: A single-nucleus transcriptomic atlas of the adult fruit fly. Science (1979) 375. 10.1126/science.abk2432.

44. Morin, X., Daneman, R., Zavortink, M., and Chia, W. A protein trap strategy to detect GFP-tagged proteins expressed from their endogenous loci in Drosophila.

45. Buszczak, M., Paterno, S., Lighthouse, D., Bachman, J., Planck, J., Owen, S., Skora, A.D., Nystul, T.G., Ohlstein, B., Allen, A., et al. (2007). The carnegie protein trap library: A versatile tool for drosophila developmental studies. Genetics 175, 1505–1531. 10.1534/genetics.106.065961.

46. Mao, Z., and Davis, R.L. (2009). Eight different types of dopaminergic neurons innervate the Drosophila mushroom body neuropil: Anatomical and physiological heterogeneity. Front Neural Circuits 3, 5. 10.3389/NEURO.04.005.2009/BIBTEX.

47. Voet, M. Van Der, Harich, B., Franke, B., and Schenck, A. (2016). ADHD-associated dopamine transporter, latrophilin and neurofibromin share a dopamine-related locomotor signature in Drosophila. Mol Psychiatry 21, 565–573. 10.1038/mp.2015.55.

48. Kume, K., Kume, S., Park, S.K., Hirsh, J., and Jackson, F.R. (2005). Dopamine is a regulator of arousal in the fruit fly. Journal of Neuroscience 25, 7377–7384. 10.1523/JNEUROSCI.2048-05.2005.

49. Nall, A.H., Shakhmantsir, I., Cichewicz, K., Birman, S., Hirsh, J., and Sehgal, A. (2016). Caffeine promotes wakefulness via dopamine signaling in Drosophila. Sci Rep 6. 10.1038/srep20938.

50. Ni, J.Q., Markstein, M., Binari, R., Pfeiffer, B., Liu, L.P., Villalta, C., Booker, M., Perkins, L., and Perrimon, N. (2007). Vector and parameters for targeted transgenic RNA interference in Drosophila melanogaster. Nature Methods 2008 5:1 5, 49–51. 10.1038/nmeth1146.

51. McGary, K.L., Park, T.J., Woods, J.O., Cha, H.J., Wallingford, J.B., and Marcotte, E.M. (2010). Systematic discovery of nonobvious human disease models through orthologous phenotypes. Proc Natl Acad Sci U S A 107, 6544–6549. 10.1073/pnas.0910200107.

52. Nakashima, A., Ohnuma, S., Kodani, Y., Kaneko, Y.S., Nagasaki, H., Nagatsu, T., and Ota, A. (2016). Inhibition of deubiquitinating activity of USP14 decreases tyrosine hydroxylase phosphorylated at Ser19 in PC12D cells. Biochem Biophys Res Commun 472, 598–602. 10.1016/J.BBRC.2016.03.022.

53. Kawahata, I., Ohtaku, S., Tomioka, Y., Ichinose, H., and Yamakuni, T. (2015). Dopamine or biopterin deficiency potentiates phosphorylation at (40)Ser and ubiquitination of tyrosine hydroxylase to be degraded by the ubiquitin proteasome system. Biochem Biophys Res Commun 465, 53–58. 10.1016/J.BBRC.2015.07.125.

54. Birman, S., Morgan, B., Anzivino, M., and Hirsh, J. (1994). A novel and major isoform of tyrosine hydroxylase in Drosophila is generated by alternative RNA processing. Journal of Biological Chemistry 269, 26559–26567. 10.1016/S0021-9258(18)47231-6.

55. Vié, A., Cigna, M., Toci, R., and Birman, S. (1999). Differential regulation of Drosophila tyrosine hydroxylase isoforms by dopamine binding and cAMP-dependent phosphorylation. J Biol Chem 274, 16788–16795. 10.1074/JBC.274.24.16788.

56. Gervasi, N.M., Scott, S.S., Aschrafi, A., Gale, J., Vohra, S.N., Macgibeny, M.A., Kar, A.N., Gioio, A.E., and Kaplan, B.B. (2016). The local expression and trafficking of tyrosine hydroxylase mRNA in the axons of sympathetic neurons. RNA 22, 883–895. 10.1261/RNA.053272.115.

57. Freeman, A., Pranski, E., Miller, R.D., Radmard, S., Bernhard, D., Jinnah, H.A., Betarbet, R., Rye, D.B., and Sanyal, S. (2012). Sleep fragmentation and motor restlessness in a Drosophila model of Restless Legs Syndrome. Current Biology 22, 1142. 10.1016/J.CUB.2012.04.027.

58. Chaudhuri, A., Bowling, K., Funderburk, C., Lawal, H., Inamdar, A., Wang, Z., and O’Donnell, J.M. (2007). Interaction of Genetic and Environmental Factors in a Drosophila Parkinsonism Model. The Journal of Neuroscience 27, 2457. 10.1523/JNEUROSCI.4239-06.2007.

59. Wang, Z., Ferdousy, F., Lawal, H., Huang, Z., Daigle, J.G., Izevbaye, I., Doherty, O., Thomas, J., Stathakis, D.G., and O’Donnell, J.M. (2011). Catecholamines up integrates dopamine synthesis and synaptic trafficking. J Neurochem 119, 1294–1305. 10.1111/J.1471-4159.2011.07517.X.

60. Zhang, Y.Q., Friedman, D.B., Wang, Z., Woodruff, E., Pan, L., O’Donnell, J., and Broadie, K. (2005). Protein expression profiling of the Drosophila fragile X mutant brain reveals up-regulation of monoamine synthesis. Molecular and Cellular Proteomics 4, 278–290. 10.1074/mcp.M400174-MCP200.

61. Wicker-Thomas, C., and Hamann, M. (2008). Interaction of dopamine, female pheromones, locomotion and sex behavior in Drosophila melanogaster. J Insect Physiol 54, 1423–1431. 10.1016/j.jinsphys.2008.08.005.

62. Takahashi, A. (2013). Pigmentation and behavior: potential association through pleiotropic genes in Drosophila.

63. Letizia, A., Bario, R., and Campuzano, S. (2007). Antagonistic and cooperative actions of the EGFR and Dpp pathways on the iroquois genes regulate Drosophila mesothorax specification and patterning. Development 134, 1337–1346. 10.1242/DEV.02823.

64. Lu, J., Wang, Y., Wang, X., Wang, D., Pflugfelder, G.O., and Shen, J. (2022). The Tbx6 Transcription Factor Dorsocross Mediates Dpp Signaling to Regulate Drosophila Thorax Closure. Int J Mol Sci 23. 10.3390/IJMS23094543.

65. Zeitlinger, J., and Bohmann, D. (1999). Thorax closure in Drosophila: involvement of Fos and the JNK pathway. Development 126, 3947–3956. 10.1242/DEV.126.17.3947.

66. Massey, J.H., Akiyama, N., Bien, T., Dreisewerd, K., Wittkopp, P.J., Yew, J.Y., and Takahashi, A. (2019). Pleiotropic Effects of ebony and tan on Pigmentation and Cuticular Hydrocarbon Composition in Drosophila melanogaster. Front Physiol 10, 518. 10.3389/FPHYS.2019.00518/BIBTEX.

67. Zhang, S., Wang, R., and Wang, G. (2019). Impact of Dopamine Oxidation on Dopaminergic Neurodegeneration. ACS Chem Neurosci 10, 945–953. 10.1021/ACSCHEMNEURO.8B00454/ASSET/IMAGES/MEDIUM/CN-2018-004545_0005.GIF.

68. Davla, S., Artiushin, G., Li, Y., Chitsaz, D., Li, S., Sehgal, A., and van Meyel, D.J. (2020). AANAT1 functions in astrocytes to regulate sleep homeostasis. Elife 9, 1–48. 10.7554/ELIFE.53994.

69. DeAngelis, M.W., McGhie, E.W., Coolon, J.D., and Johnson, R.I. (2020). Mask, a component of the Hippo pathway, is required for Drosophila eye morphogenesis. Dev Biol 464, 53–70. 10.1016/j.ydbio.2020.05.002.

70. Zhu, M., Zhang, S., Tian, X., and Wu, C. (2017). Mask mitigates MAPT- and FUS-induced degeneration by enhancing autophagy through lysosomal acidification. Autophagy 13, 1924–1938. 10.1080/15548627.2017.1362524.

71. Martinez, D., Zhu, M., Guidry, J.J., Majeste, N., Mao, H., Yanofsky, S.T., Tian, X., and Wu, C. (2021). Mask, the Drosophila ankyrin repeat and KH domain-containing protein, affects microtubule stability. J Cell Sci 134. 10.1242/jcs.258512.

72. Dubowy, C., and Sehgal, A. (2017). Circadian rhythms and sleep in Drosophila melanogaster. Genetics 205, 1373–1397. 10.1534/genetics.115.185157.

73. Cichewicz, K., Garren, E.J., Adiele, C., Aso, Y., Wang, Z., Wu, M., Birman, S., Rubin, G.M., and Hirsh, J. (2017). A new brain dopamine-deficient Drosophila and its pharmacological and genetic rescue. Genes Brain Behav 16, 394–403. 10.1111/GBB.12353.

74. Sidor, C.M., Brain, R., and Thompson, B.J. (2013). Mask proteins are cofactors of Yorkie/YAP in the Hippo pathway. Curr Biol 23, 223–228. 10.1016/J.CUB.2012.11.061.

75. Sansores-Garcia, L., Atkins, M., Moya, I.M., Shahmoradgoli, M., Tao, C., Mills, G.B., and Halder, G. (2013). Mask is required for the activity of the Hippo pathway effector Yki/YAP. Curr Biol 23, 229–235. 10.1016/J.CUB.2012.12.033.

76. Smith, R.K., Carroll, P.M., Allard, J.D., and Simon, M.A. (2002). MASK, a large ankyrin repeat and KH domain-containing protein involved in Drosophila receptor tyrosine kinase signaling. Development 129, 71–82. 10.1242/DEV.129.1.71.

77. Zhu, M., Li, X., Tian, X., and Wu, C. (2015). Mask loss-of-function rescues mitochondrial impairment and muscle degeneration of Drosophila pink1 and parkin mutants. Hum Mol Genet 24, 3272–3285. 10.1093/HMG/DDV081.

78. Lusk, J.B., Lam, V.Y.M., and Tolwinski, N.S. (2017). Epidermal Growth Factor Pathway Signaling in Drosophila Embryogenesis: Tools for Understanding Cancer. Cancers (Basel) 9. 10.3390/CANCERS9020016.

79. Oh, H., and Irvine, K.D. (2010). Yorkie: the final destination of Hippo signaling. Trends Cell Biol 20, 410. 10.1016/J.TCB.2010.04.005.

80. Sen, A., Kalvakuri, S., Bodmer, R., and Cox, R.T. (2015). Clueless, a protein required for mitochondrial function, interacts with the PINK1-Parkin complex in Drosophila. Dis Model Mech 8, 577–589. 10.1242/DMM.019208.

81. Wang, Z.H., Clark, C., and Geisbrecht, E.R. (2016). Drosophila clueless is involved in Parkin-dependent mitophagy by promoting VCP-mediated Marf degradation. Hum Mol Genet 25, 1946–1964. 10.1093/HMG/DDW067.

82. Sen, A., and Cox, R.T. (2016). Clueless is a conserved ribonucleoprotein that binds the ribosome at the mitochondrial outer membrane. Biol Open 5, 195–203. 10.1242/BIO.015313/-/DC1.

83. Inagaki, H.K., De-Leon, S.B.-T., Wong, A.M., Jagadish, S., Ishimoto, H., Barnea, G., Kitamoto, T., Axel, R., and Anderson, D.J. (2012). Visualizing Neuromodulation In Vivo: TANGO-Mapping of Dopamine Signaling Reveals Appeptite Control of Sugar Sensing. Cell 148, 583. 10.1016/J.CELL.2011.12.022.

84. Sun, F., Zhou, J., Dai, B., Qian, T., Zeng, J., Li, X., Zhuo, Y., Zhang, Y., Wang, Y., Qian, C., et al. (2020). Next-generation GRAB sensors for monitoring dopaminergic activity in vivo. Nat Methods 17, 1156. 10.1038/S41592-020-00981-9.

85. Kasture, A.S., Hummel, T., Sucic, S., and Freissmuth, M. (2018). Big Lessons from Tiny Flies: Drosophila melanogaster as a Model to Explore Dysfunction of Dopaminergic and Serotonergic Neurotransmitter Systems. Int J Mol Sci 19. 10.3390/IJMS19061788.

86. Calleja, M., Herranz, H., Estella, C., Casal, J., Lawrence, P., Simpson, P., and Morata, G. (2000). Generation of medial and lateral dorsal body domains by the pannier gene of Drosophila. Development 127, 3971–3980. 10.1242/DEV.127.18.3971.

87. Kennerdell, J.R., and Carthew, R.W. (2000). Heritable gene silencing in Drosophila using double-stranded RNA. Nat Biotechnol 18, 896–898. 10.1038/78531.

88. Yamamoto, S., Charng, W.L., Rana, N.A., Kakuda, S., Jaiswal, M., Bayat, V., Xiong, B., Zhang, K., Sandoval, H., David, G., et al. (2012). A mutation in EGF repeat-8 of notch discriminates between serrate/jagged and delta family ligands. Science (1979) 338, 1229–1232. 10.1126/science.1228745.

89. Marcogliese, P.C., Deal, S.L., Andrews, J., Harnish, J.M., Bhavana, V.H., Graves, H.K., Jangam, S., Luo, X., Liu, N., Bei, D., et al. (2022). Drosophila functional screening of de novo variants in autism uncovers damaging variants and facilitates discovery of rare neurodevelopmental diseases. Cell Rep 38. 10.1016/j.celrep.2022.110517.

90. Snel, B., Lehmann, G., Bork, P., and Huynen, M.A. (2000). STRING: a web-server to retrieve and display the repeatedly occurring neighbourhood of a gene.

91. Davie, K., Janssens, J., Koldere, D., De Waegeneer, M., Pech, U., Kreft, Ł., Aibar, S., Makhzami, S., Christiaens, V., Bravo González-Blas, C., et al. (2018). A Single-Cell Transcriptome Atlas of the Aging Drosophila Brain. Cell 174, 982–998.e20. 10.1016/j.cell.2018.05.057.

92. Szklarczyk, D., Gable, A.L., Nastou, K.C., Lyon, D., Kirsch, R., Pyysalo, S., Doncheva, N.T., Legeay, M., Fang, T., Bork, P., et al. (2021). The STRING database in 2021: Customizable protein-protein networks, and functional characterization of user-uploaded gene/measurement sets. Nucleic Acids Res 49, D605–D612. 10.1093/nar/gkaa1074.

93. Szklarczyk, D., Kirsch, R., Koutrouli, M., Nastou, K., Mehryary, F., Hachilif, R., Gable, A.L., Fang, T., Doncheva, N.T., Pyysalo, S., et al. (2023). The STRING database in 2023: protein-protein association networks and functional enrichment analyses for any sequenced genome of interest. Nucleic Acids Res 51, D638–D646. 10.1093/nar/gkac1000.

94. Hu, Y., Flockhart, I., Vinayagam, A., Bergwitz, C., Berger, B., Perrimon, N., and Mohr, S.E. (2011). An integrative approach to ortholog prediction for disease-focused and other functional studies. BMC Bioinformatics 12. 10.1186/1471-2105-12-357.

95. Hamosh, A., Amberger, J.S., Bocchini, C., Scott, A.F., and Rasmussen, S.A. (2021). Online Mendelian Inheritance in Man (OMIM®): Victor McKusick’s magnum opus. Am J Med Genet A 185, 3259–3265. 10.1002/ajmg.a.62407.

96. Abrahams, B.S., Arking, D.E., Campbell, D.B., Mefford, H.C., Morrow, E.M., Weiss, L.A., Menashe, I., Wadkins, T., Banerjee-Basu, S., and Packer, A. (2013). SFARI Gene 2.0: A community-driven knowledgebase for the autism spectrum disorders (ASDs). Mol Autism 4. 10.1186/2040-2392-4-36.

97. Cichewicz, K., Garren, E.J., Adiele, C., Aso, Y., Wang, Z., Wu, M., Birman, S., Rubin, G.M., and Hirsh, J. (2017). A new brain dopamine-deficient Drosophila and its pharmacological and genetic rescue. Genes Brain Behav 16, 394–403. 10.1111/gbb.12353.

98. Hardie, S.L., and Hirsh, J. (2006). An improved method for the separation and detection of biogenic amines in adult Drosophila brain extracts by high performance liquid chromatography. J Neurosci Methods 153, 243–249. 10.1016/j.jneumeth.2005.11.001.

