## Supplemental Figures - All for "RNAi-based screen for pigmentation in *Drosophila melanogaster* reveals regulators of brain dopamine and sleep"

### Supplemental Figure 1

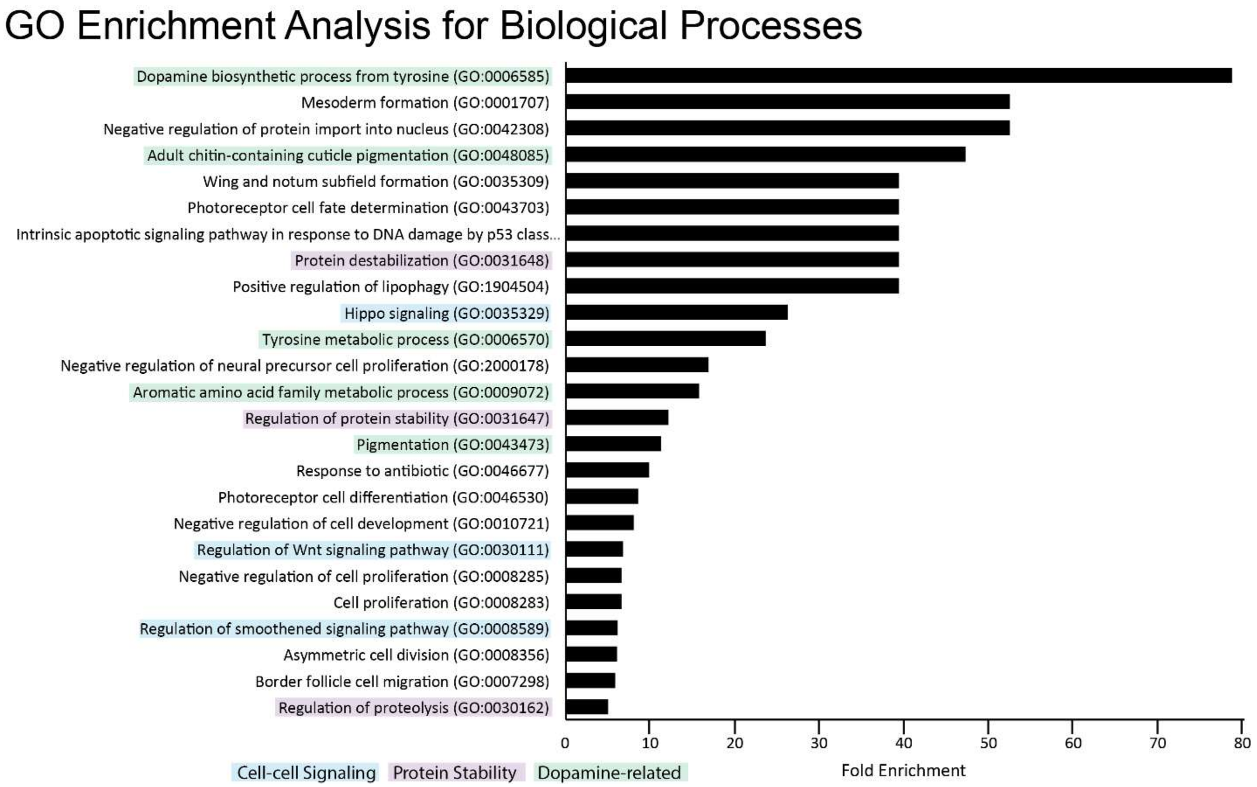


**S1 Fig. GO term enrichment analysis performed on the pigmentation screen hits.** GO term analysis on the 153 pigmentation screen hits. Highlighted are terms in related areas of biology: cell-cell signaling related terms are in blue, protein stability related terms are in purple, and dopamine related processes or pathways are highlighted in green. See materials and methods for detailed analysis information.

### Supplemental Figure 2

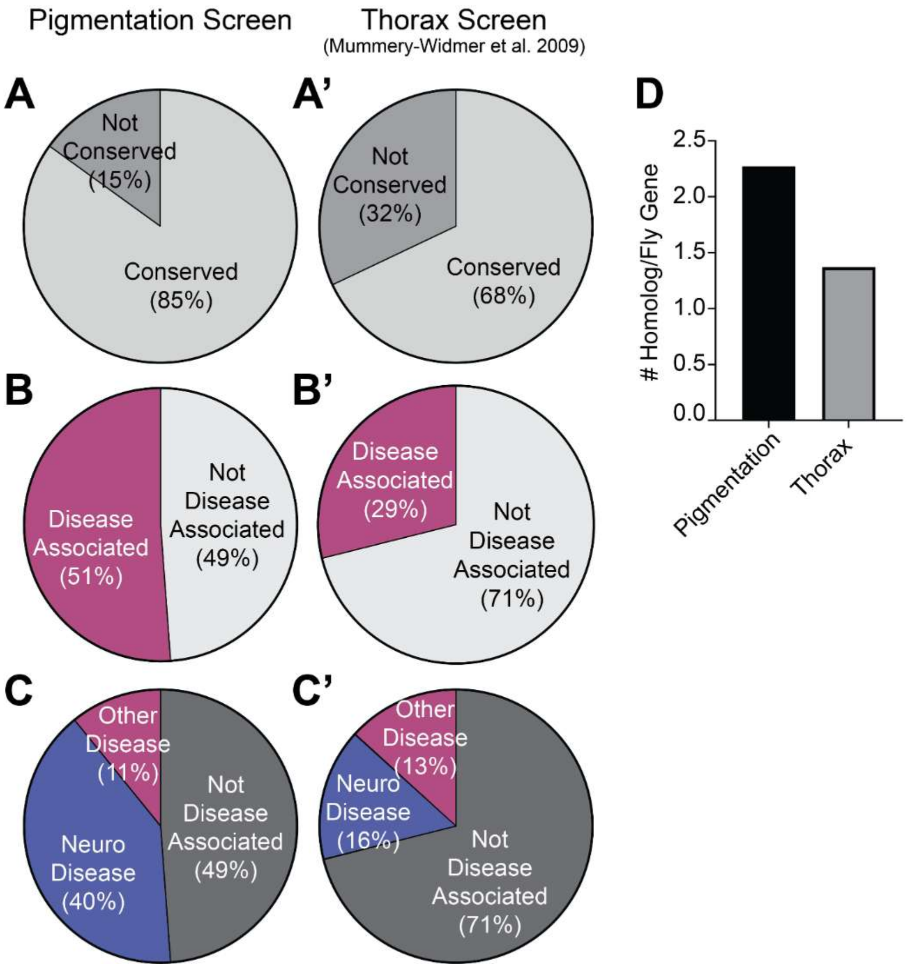


**S2 Fig. Comparison between genes identified as hits from our pigmentation screen to those tested in the original thorax screen (Mummery-Widmer et al., 2009).** (A-A') Pie charts comparing number of conserved genes. (B-B') Pie charts comparing number of fly genes that have a homolog that is associated with a disease. (C-C') Pie charts comparing the number of fly genes that have a homology associated with a neurological disease. (D) Bar graph showing the average number of homologs for the genes identified in our screen and the original screen.

### Supplemental Figure 3

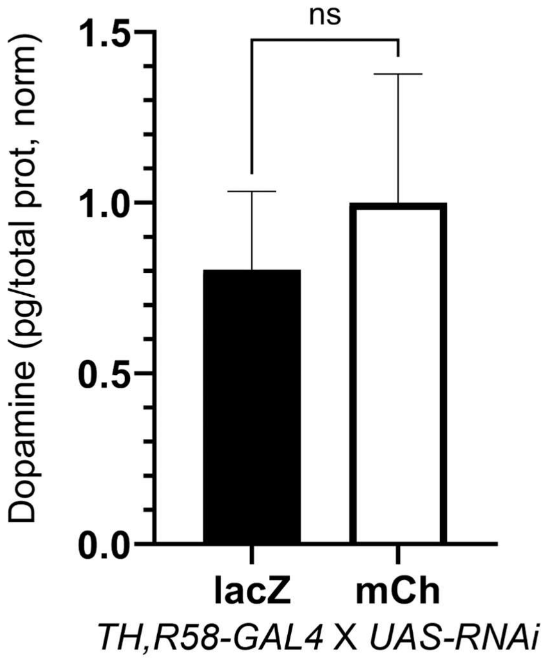


**S3 Fig. Comparison of dopamine levels between lacZ RNAi and mCh RNAi lines in the fly head.** Both lacZ-RNAi and mCherry(mCh)-RNAi lines are considered as ‘control’ lines for different RNAi collections. Statistical analysis using t-test. Error bars show SD.

### Supplemental Figure 4

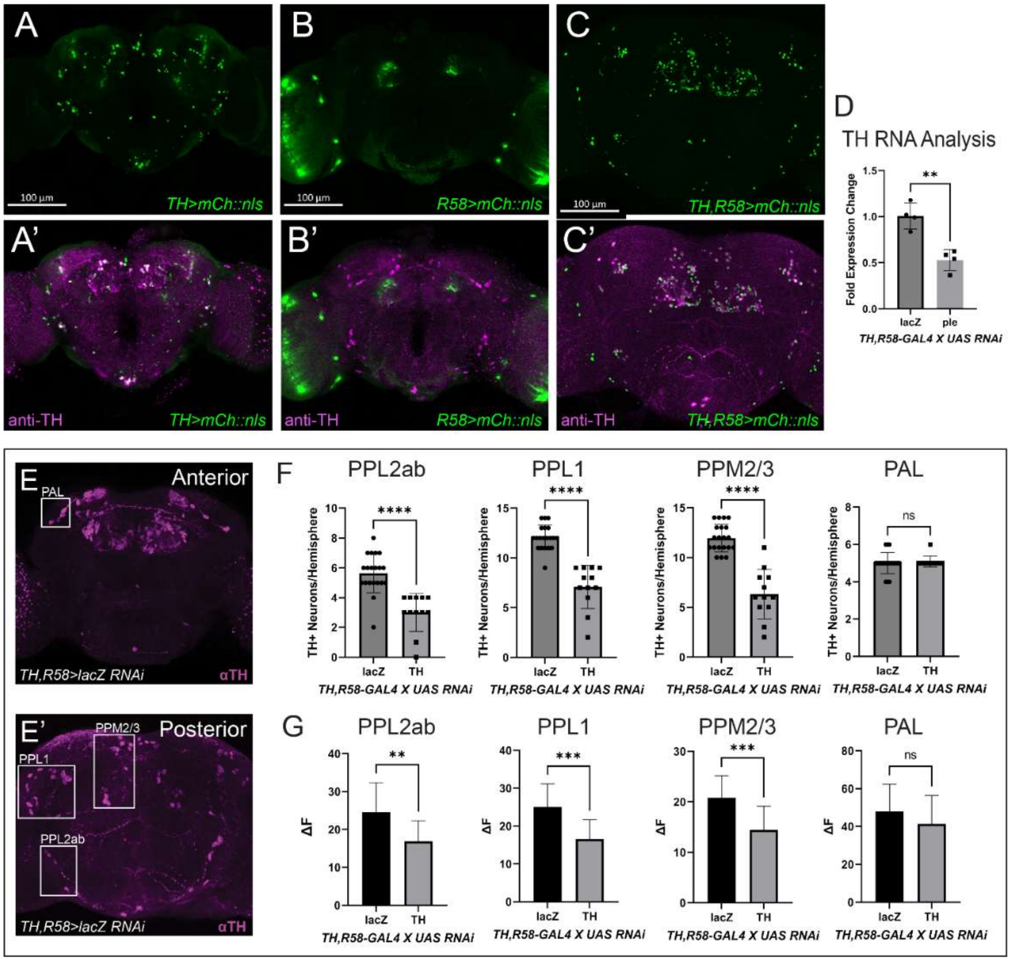


**S4 Fig. Validation of recombined GAL4 lines used for dopamine measurement in the brain.** (A-A') Expression of well-characterized *TH-GAL4* (i.e. *ple-GAL4*) in the adult brain shows overlap with most dopaminergic neurons, except for those in the PAM cluster of DANs. (B-B') *R58-GAL4* is expressed in most DANs in the PAM cluster. (C-C') Recombined *TH,R58-GAL4* is expressed in 90-95% of DANs including most of the PAM cluster. (D) TH mRNA levels upon TH knockdown using recombined GAL4 line (E-E') Dopaminergic neurons labeled with TH antibody in the anterior (E) and posterior (E') of the brain. Specific dopaminergic clusters highlighted. (F-G) Quantification of TH+ positive neurons (F) and total TH immunofluorescence (G) upon knockdown of TH with the recombined GAL4 line. **=p<0.01, ***=p<0.001, ****=p<0.0001. Error bars represent SD. Scale bars represent 100 µm.

### Supplemental Figure 5

**A**

| Gene | DANs # | % DANs |
|---|---|---|
| DAT | 317 | 98.8 |
| ple | 287 | 89.4 |
| mask | 113 | 35.2 |
| Bsg | 89 | 27.7 |
| Dpy-30L1 | 65 | 20.2 |
| ClC-c | 46 | 14.3 |
| clu | 36 | 11.2 |
| Membrin | 28 | 8.7 |
| CG43689 | 17 | 5.3 |
| rk | 13 | 4.0 |
| Sos | 12 | 3.7 |
| PIG-A | 9 | 2.8 |
| Spn38F | 0 | 0 |

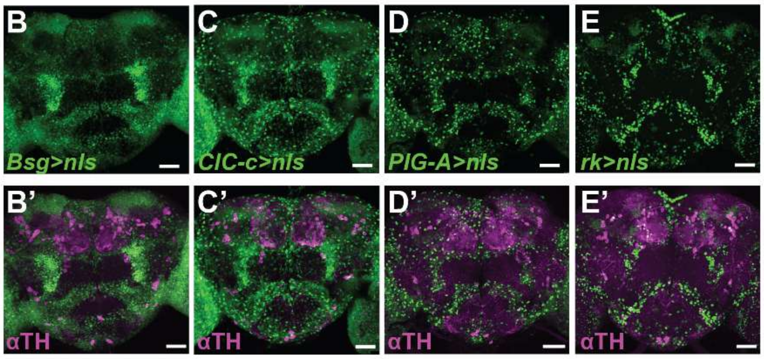


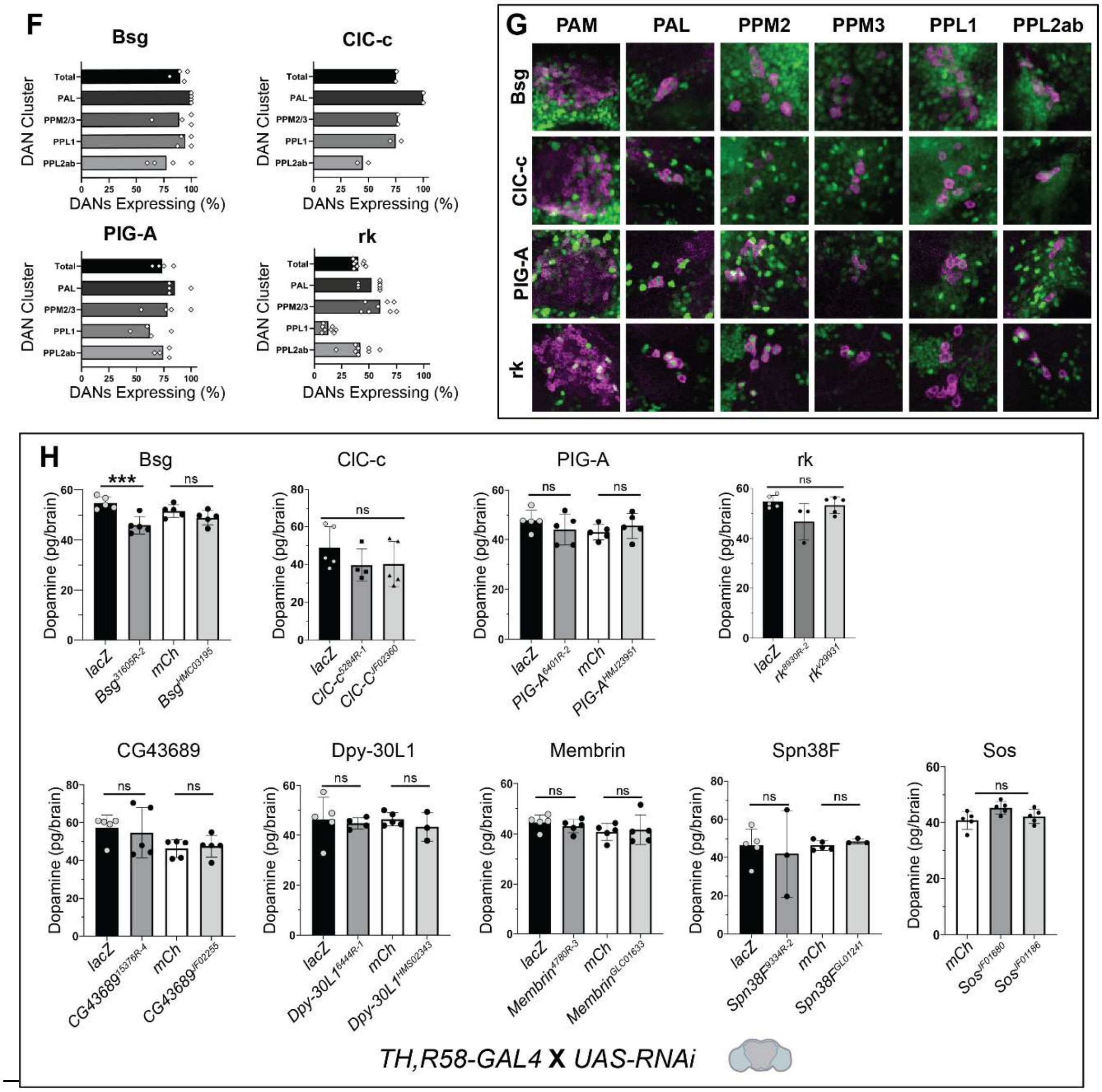

**S5 Fig. Expression analysis and brain HPLC measurements of additional prioritized genes.** (A) Quantification of overlap between the given gene and DANs. DANs defined based on DAN cluster identified in single-cell sequencing from Davie et al. (2018). (B-E) Adult brains for *Bsg*$^{T2A}$-*GAL4*, *ClC-c*$^{T2A}$-*GAL4*, *PIG-A*$^{T2A}$-*GAL4*, and *rk*$^{T2A}$-*GAL4* driving *UAS-mCh::nls* (nuclear localized mCherry). (B'-E') Overlap with DANs as labelled with anti-TH (magenta). (F) Quantification of TH+ neurons with nls expression from *Bsg*$^{T2A}$*->mCh::nls*, *ClC-c*$^{T2A}$*>mCh::nls*, *PIG-A*$^{T2A}$*>mCh::nls*, and *rk*$^{T2A}$*>mCh::nls* for the given DAN clusters. (G) Representative images of the expression of *Bsg*$^{T2A}$*->mCh::nls*, *ClC-c*$^{T2A}$*>mCh::nls*, *PIG-A*$^{T2A}$*>mCh::nls*, and *rk*$^{T2A}$*>mCh::nls* for the given DAN clusters (H) Dopamine measured from dissected adult brains using HPLC upon knockdown of *Bsg*, *ClC-c*, *PIG-A*, *rk*, *CG43689*, *Dpy-30L1*, *Membrin*, *Spn38F*, *Sos* using a recombined dopaminergic GAL4 driver (*TH, R58-GAL4*).

### Supplemental Figure 6

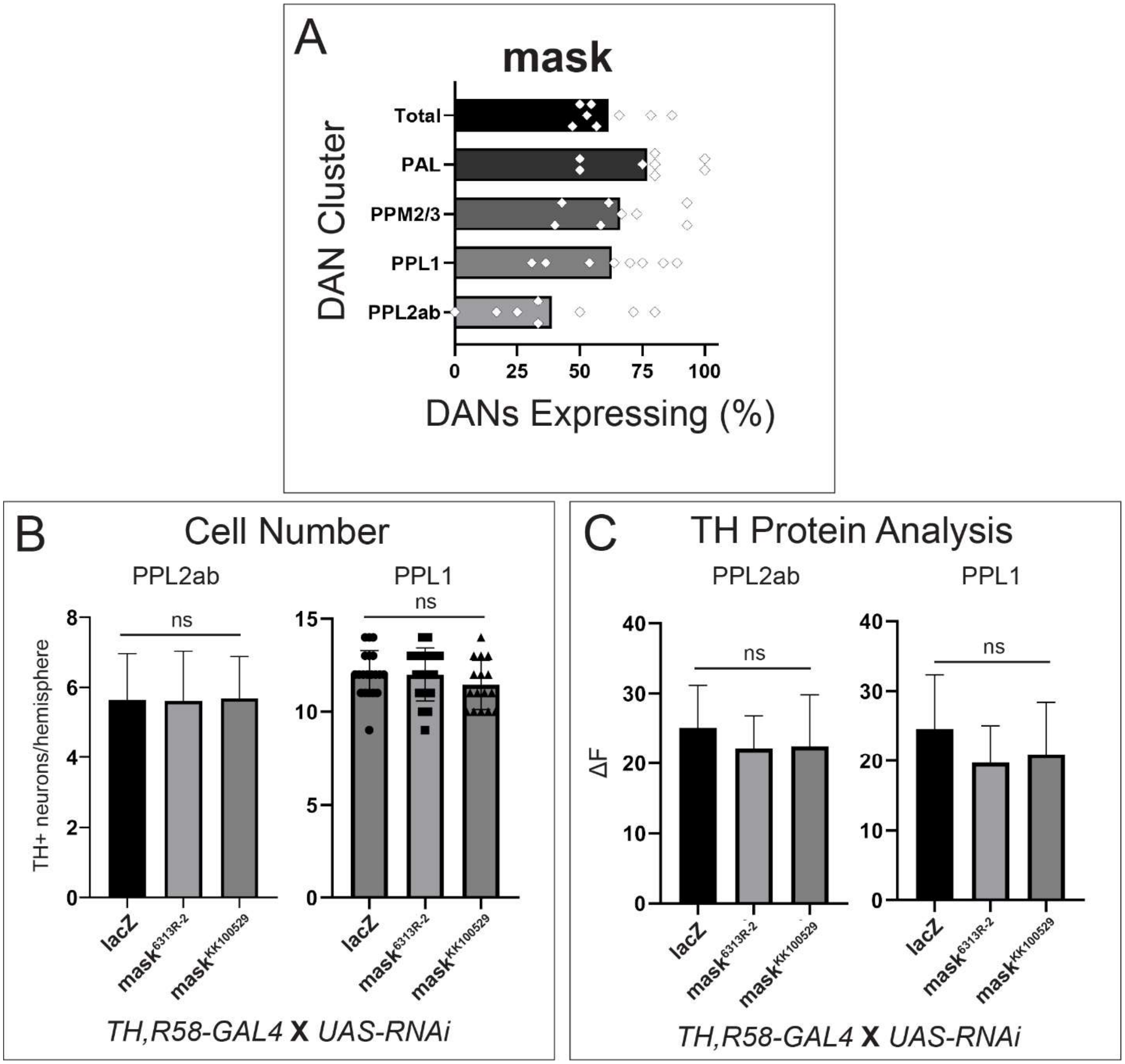


**S6 Fig. Additional data from brain studies on mask – co-expression with TH and quantification of TH positive neurons upon mask knockdown.** (A) Quantification of TH+ neurons with nls expression from *mask*$^{T2A}$*>mCh::nls* for the given DAN clusters. (B) Quantification of TH positive (TH+) cell number upon *TH* knockdown in given clusters of DANs. (C) Quantification of TH mRNA level upon *TH* knockdown using qRT-PCR. Ordinary one-way ANOVAs performed. Error bars represent SD.

### Supplemental Figure 7

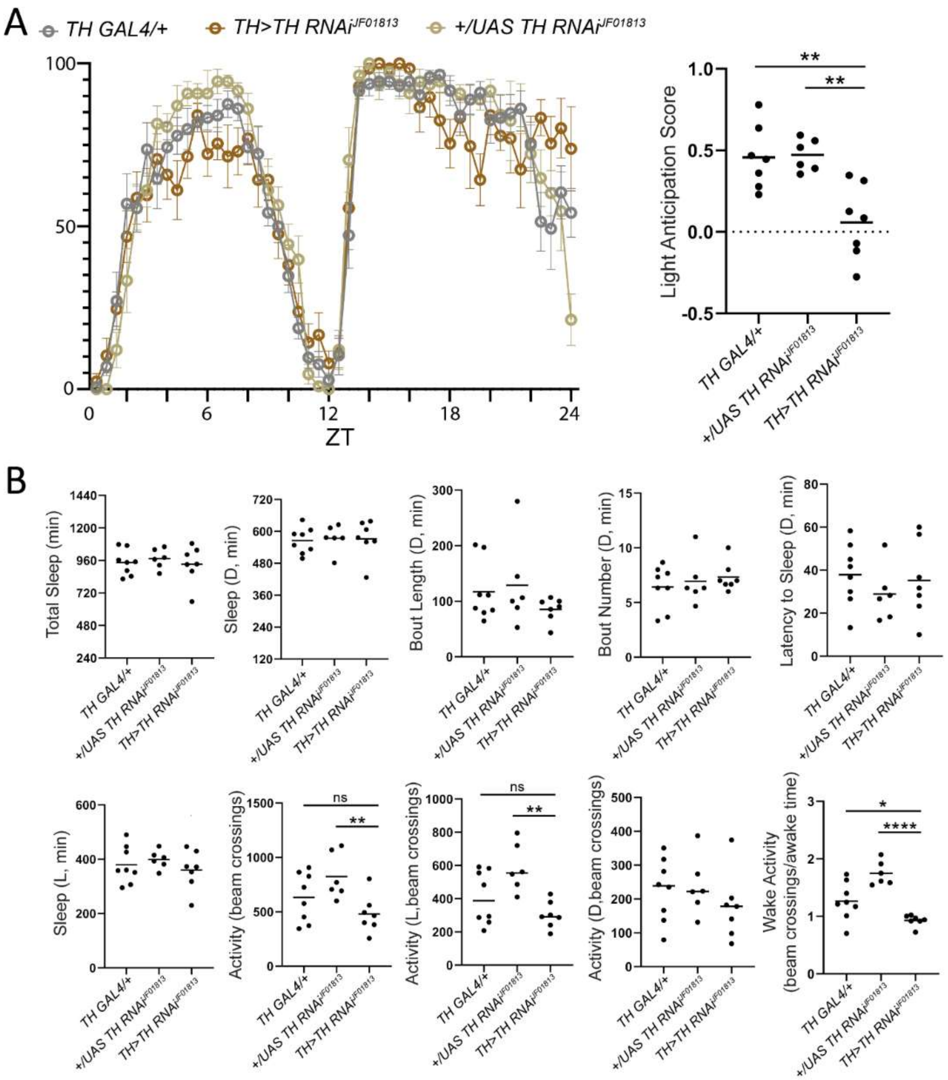


**S7 Fig. Sleep behavior analysis upon TH GAL4 driven TH knockdown.** (A) 24-hr sleep behavior (binned in 30-minute intervals) from TH>TH RNAi compared to controls shows increased sleep in the last two hours of the dark period. Quantification of light anticipation highlights effect on that last two hours before light onset. (B) Otherwise, sleep is not affected in TH>TH RNAi with the exception of Wake Activity, which is reduced in the TH>TH RNAi experiment. Ordinary one-way ANOVAs performed with Dunnett's multiple comparisons for those with significant differences. *=p<0.05, **=p<0.01, ****=p<0.0001. Error bars represent SD.

### Supplemental Figure 8

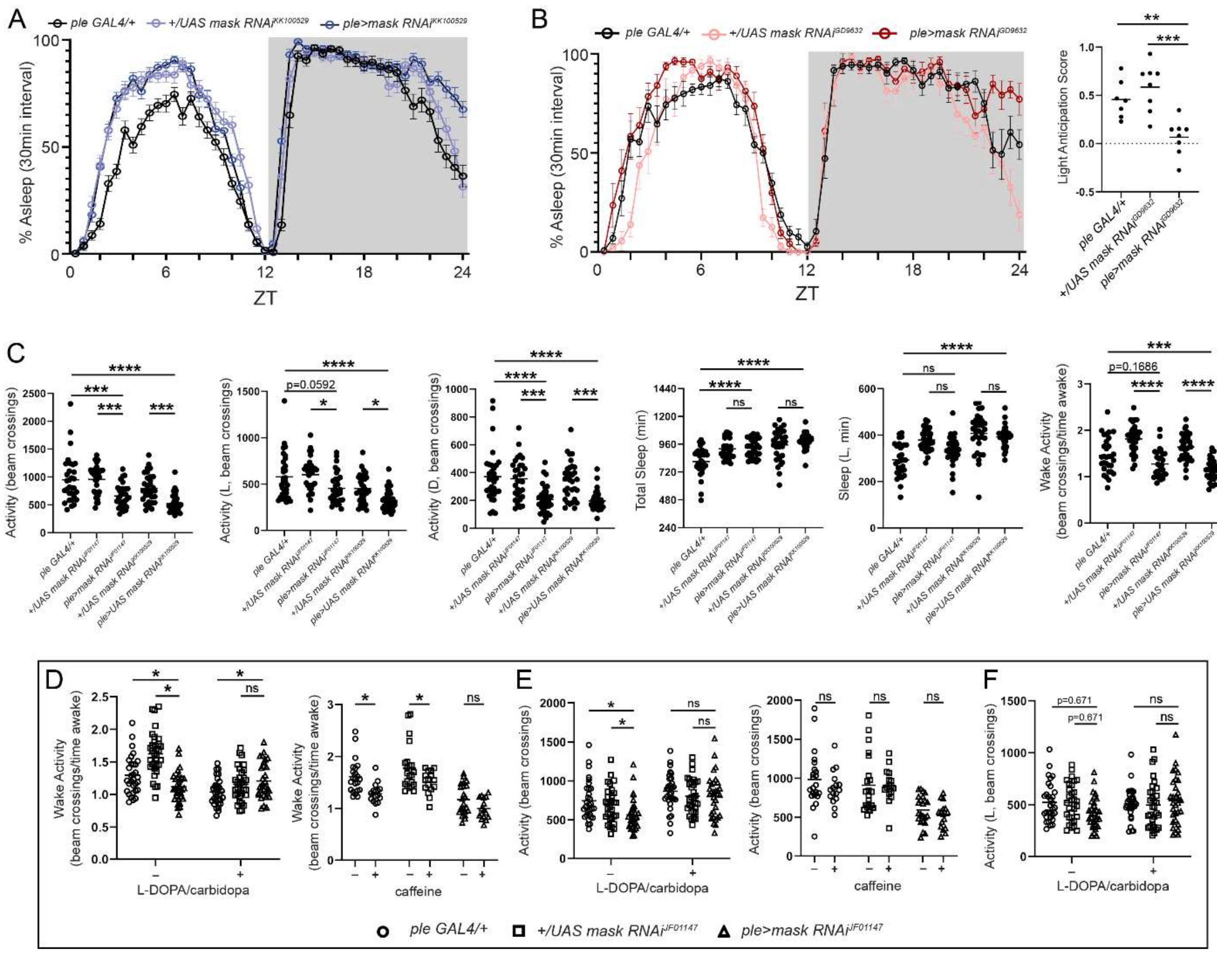


**S8 Fig. Additional sleep analysis from TH-GAL4 driven *mask* knockdown.** (A) Additional RNAi lines tested for sleep phenotypes showed increased sleep before light onset, also known as reduced light anticipation. (A') Quantification for an additional RNAi line. Other line quantification found in Fig 6. (B) Additional Drosophila Activity Monitor parameters measured from TH>mask RNAi experiments show reduced activity and reduced wake activity, with no significant effect on total sleep or sleep during the 12-hr light period. (C) Wake activity can also be rescued with L-DOPA and caffeine effects on wake activity are not seen upon mask knockdown. (D-E) Total activity effects are also rescued by L-DOPA, but there are no overall effects on activity upon caffeine. There are also no changes in activity during the light period upon mask knockdown with or without L-DOPA.

### Supplemental Figure 9

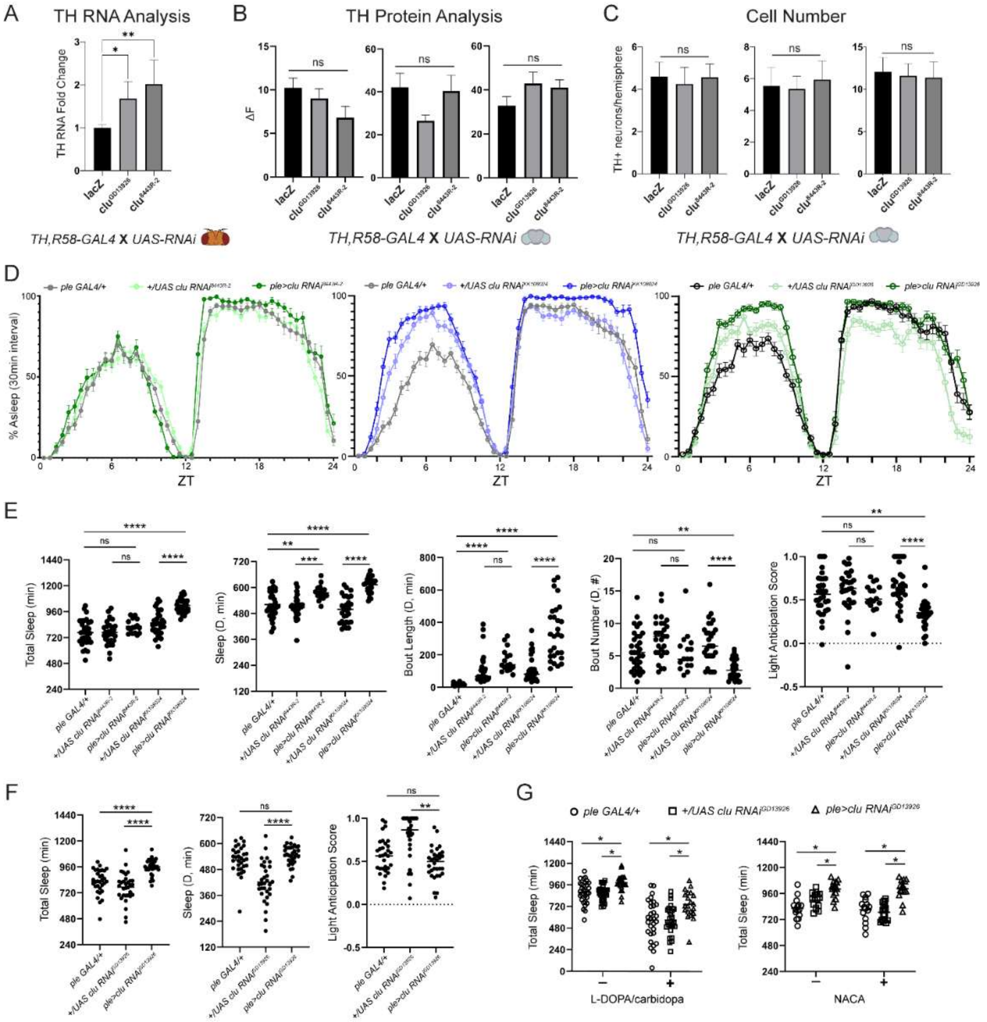


**S9 Fig. Sleep behavior analysis of *clueless* knockdown highlights a different pathway than mask.** (A) qRT-PCR analysis on TH mRNA levels shows that clueless knockdown leads to increases in TH mRNA. (B-C) Image analysis using TH immunofluorescent antibody staining shows no effect on TH protein levels (B) and no change in TH+ cell numbers (C) upon *clueless* knockdown with TH-GAL4. (D) 24-hr sleep behavior graphs (binned in 30-minute intervals) show inconsistency for time of day when sleep is increase. (E-F) Quantification shows that there is a consistent increase in sleep upon clueless knockdown, but no effect on light anticipation. (G) The effect on total sleep upon *clueless* knockdown is not rescued by feeding flies L-DOPA or the antioxidant NACA.

### Supplemental Figure 10

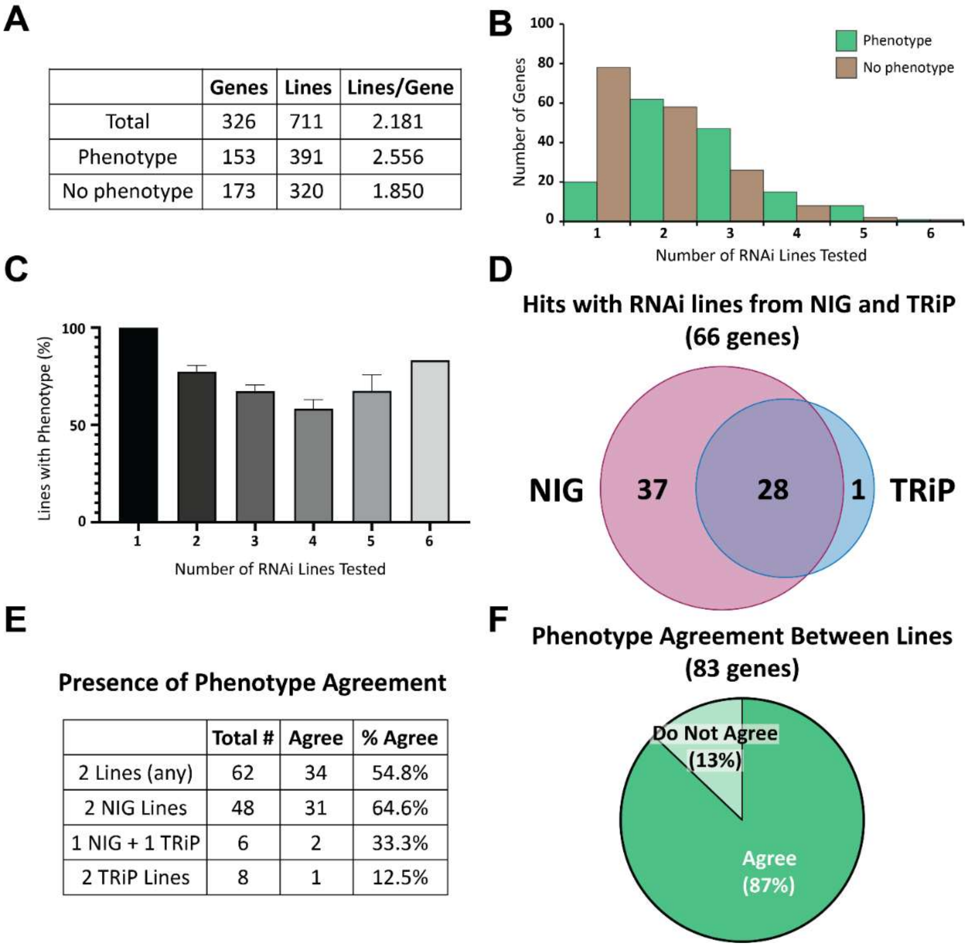


| | Genes | Lines | Lines/Gene |
|---|---|---|---|
| Total | 326 | 711 | 2.181 |
| Phenotype | 153 | 391 | 2.556 |
| No phenotype | 173 | 320 | 1.850 |

| | Total # | Agree | % Agree |
|---|---|---|---|
| 2 Lines (any) | 62 | 34 | 54.8% |
| 2 NIG Lines | 48 | 31 | 64.6% |
| 1 NIG + 1 TRiP | 6 | 2 | 33.3% |
| 2 TRiP Lines | 8 | 1 | 12.5% |

**S10 Fig. Comparison of RNAi lines from the pigmentation screen** (A) Number of lines tested for a given gene. (B) Histogram comparing number of lines tested for those that showed a phenotype and those that did not. (C) Bar graph depicting the number of lines that showed a phenotype by the number of lines tested. (i.e. for those where two lines were tested ~77% of them showed a phenotype). (D) Venn diagram comparing genes with both NIG and TRiP RNAi lines showed that NIG lines were more likely to give a phenotype. (E) Comparison for those genes that only have two lines tested showed agreement (in terms of phenotype or no phenotype) was ~55% across all genes, but most of this agreement comes from NIG lines tested. (F) Pie chart showing phenotypic agreement (i.e. all 'pale' or all 'dark') for the 83 genes where multiple lines had a phenotype. Error bars represent standard error of the mean.

### Supplemental Figure 11

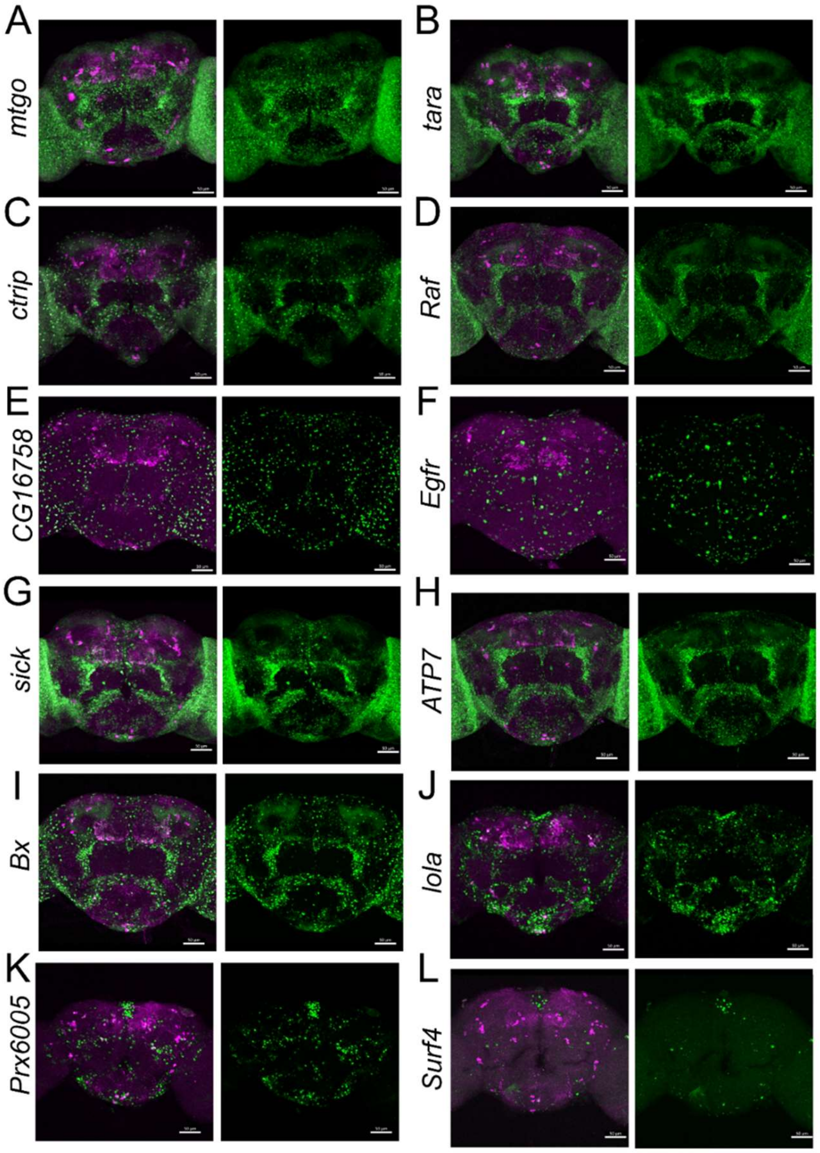


**S11 Fig. Expression analysis for other prioritized candidate genes that did not show any significant change in total head dopamine**. Expression pattern of *T2A-GAL4* lines available for *mtgo* (A), *tara* (B), *ctrip* (C), *Raf* (D), *CG16758* (E), *Egfr* (F), *sick* (G), *ATP7* (H), *Bx* (I), *lola* (J), *Prx6005* (K), *Surf4* (L). Scale bar represents 50 μm.
